# Adrenergic C1 neurons enhance anxiety via projections to PAG

**DOI:** 10.1101/2024.09.11.612440

**Authors:** Carlos Fernández-Peña, Rachel L. Pace, Lourds M. Fernando, Brittany G. Pittman, Lindsay A. Schwarz

## Abstract

Anxiety is an emotional state precipitated by the anticipation of real or potential threats. Anxiety disorders are the most prevalent psychiatric illnesses globally and increase the risk of developing comorbid conditions that negatively impact the brain and body. The etiology of anxiety disorders remains unresolved, limiting improvement of therapeutic strategies to alleviate anxiety-related symptoms with increased specificity and efficacy. Here, we applied novel intersectional tools to identify a discrete population of brainstem adrenergic neurons, named C1 cells, that promote aversion and anxiety-related behaviors via projections to the periaqueductal gray matter (PAG). While C1 cells have traditionally been implicated in modulation of autonomic processes, rabies tracing revealed that they receive input from brain areas with diverse functions. Calcium-based in vivo imaging showed that activation of C1 cells enhances excitatory responses in vlPAG, activity that is exacerbated in times of heightened stress. Furthermore, inhibition of C1 cells impedes the development of anxiety-like behaviors in response to stressful situations. Overall, these findings suggest that C1 neurons are positioned to integrate complex information from the brain and periphery for the promotion of anxiety-like behaviors.

## Introduction

Anxiety, a state of unease caused by anticipation of real or perceived threats, is associated with symptoms that encompass cognitive and autonomic functions, like shortness of breath and increased heart rate^1–3^. Anxiety disorders, including generalized anxiety disorders and panic disorders, are the most prevalent psychiatric illnesses in the world, with more than 300 million cases reported globally^4^. These diseases significantly impact quality of life and often promote development of other disorders^5,6^. Indeed, anxiety disorder patients are disproportionately affected by comorbid conditions, including cardiovascular disease, diabetes, fibromyalgia, and gastrointestinal illness^7–10^. Current treatments for anxiety disorders, such as psychotherapy and pharmacotherapy, can be effective, but often require long-term commitment from patients, as relapse is often observed when treatments are stopped^11^. Moreover, drugs used to treat anxiety are associated with adverse effects such as jitteriness, insomnia, central nervous system depression, and impaired cognitive functions, and are contraindicated for long term use^11^. Collectively, this emphasizes a need continued need for novel targets of anxiety-related therapies that function with improved specificity and long-term potency.

Despite its clinical importance, our understanding of the neurobiology of anxiety disorders remains understudied^1,12^. Studies in humans and rodents have identified several brain regions involved in anxiety, including the amygdala, prefrontal cortex, and the periaqueductal gray matter (PAG)^5,6,13–16^, but the underlying etiology and neural mechanisms by which these regions promote anxiety-related phenotypes is incomplete^17^. Among these areas, PAG is notable in that it integrates physiological functions, such as breathing and blood pressure regulation, with emotional responses and modulation of aversive and defensive behaviors^13,18–22^. PAG is subdivided into four columns: dorsal (dPAG), dorsolateral (dlPAG), lateral (lPAG) and ventrolateral (vlPAG), that are thought to preferentially process distinct behavioral responses^23,24^. For instance, while activation of dlPAG promotes escape-related behavior, activation of lPAG and vlPAG instead causes animals to preferentially freeze^18,25–29^. Meanwhile, sympathoexcitatory responses elicited by stressful stimuli (e.g. increased heart rate and breathing) are mediated by projections from lPAG and vlPAG to the rostral ventrolateral medulla (RVLM)^30^, a region located in the ventral brainstem involved in the regulation of blood pressure, respiration, and glycemia^31,32^. It has also been reported that neurons within RVLM project back to PAG^33^, introducing the possibility that RVLM itself could directly modulate anxiety-like behaviors.

RVLM contains a diversity of cell types^34^, including two subpopulations of catecholaminergic neurons called A1 and C1 cells. While both group of neurons express the enzyme dopamine-β-hydroxylase (DBH), which is required for production of norepinephrine (NE), C1 neurons also express Phenylethanolamine N-methyltransferase (PNMT), an additional enzyme in the catecholamine synthesis pathway that converts NE into epinephrine (E). Research has suggested that C1 neurons are enriched in the anterior portion of RVLM, while A1 neurons are found more caudally, with a low degree of intermingling^33^. Collectively, C1 and A1 neurons within RVLM receive inputs from a wide variety of brain regions including nucleus of the solitary tract (NTS), inferior and superior colliculi, cerebellum, and the reticular core^33^. Likewise, C1 and A1 neurons make abundant local connections and send projections to preganglionic neurons in the spinal cord and throughout the brain, innervating nuclei including amygdala, NTS, locus coeruleus (LC), and PAG^33^. Bulk activation of C1 and A1 cells induces a variety of sympathoexcitatory responses, including increases in breathing rate, blood pressure and glycemia, emphasizing their involvement in important aspects of brain-body communication^32,35–39^. Since activation of A1/C1 neurons also counteracts the disruptive physiological effects of noxious and stressful stimuli (e.g. hypoxia, hypotension), it has been suggested that C1 and A1 neurons within RVLM are critical for restoring homeostasis following stressful experiences^40,41^.

While evidence indicates that C1 and A1 neurons drive many of the physiological components of stress, it is unknown whether they play a direct role in modulating anxiety. Furthermore, as C1 and A1 cells synthesize different neurotransmitters (E versus NE), it is possible that these cell populations promote distinct actions, something that has been challenging to resolve due to the molecular similarity and close proximity of these cell groups within the RVLM. In fact, *in vivo* characterization of C1 neurons has largely relied on molecular tools that indiscriminately target all catecholaminergic cell types, making it difficult to know whether characteristics attributed to C1 cells are specific to this neuronal population.

To address this, we took advantage of recent advances in the area of intersectional molecular tools to dissociate the contributions of C1 and A1 neural circuits towards promotion of anxiety-like behaviors. Using high resolution and three-dimensional labeling strategies, we found that C1 and A1 neuronal populations are more intermingled within RVLM than anticipated. Additionally, anatomical tracing revealed that while both C1 and A1 neurons project to PAG, their axons diverge within this structure to contact distinct subregions. Combined activation of C1 and A1 neurons induces aversion and enhances anxiety-like behaviors, effects that are replicated by selective activation of C1 cells, both at the level of C1 cell bodies and through targeting of C1 axons within PAG. Monitoring endogenous C1 neuron activity reveals that they are rapidly activated in stressful situations, which in turn enhances excitatory activity in vlPAG to promote anxiety-like behaviors. Finally, inhibition of C1 neurons is anxiolytic and reduces the activity of excitatory neurons in vlPAG in response to elevated stress. Collectively, this work implicates C1 neurons as novel modulators of anxiety-induced responses via connectivity with vlPAG.

## Results

### Development of intersectional strategies for targeting C1 cells

To selectively target RVLM C1 neurons, we utilized a transgenic approach whereby expression of Cre recombinase is regulated by the *Pnmt* promoter. Although a *Pnmt^Cre^* transgenic line was previously applied to study adrenergic cells in the peripheral nervous system^42^, we found Cre expression to be highly ectopic in the brains of these mice, as indicated by the presence of a Cre-dependent reporter (tdTomato) in non-catecholaminergic cells (determined by the lack of tyrosine hydroxylase, or TH, expression) throughout the brainstem (Extended Data Fig. 1a). To overcome this issue, we generated a new transgenic mouse line, *Pnmt^2a-iCre^*, which preserves the *Pnmt* locus through insertion of a 2a-iCre sequence into the 3’ untranslated region (UTR) of the gene. We compared the specificity and efficiency of these two transgenic lines by injecting an AAV expressing Cre-dependent GFP into RVLM of *Pnmt^Cre^*and *Pnmt^2a-iCre^* mice (Extended Data Fig. 1b). Whereas non-specific AAV expression was visible within and beyond RVLM in *Pnmt^Cre^*animals, labeling was largely restricted to catecholaminergic neurons (identified via TH immunostaining) within RVLM of *Pnmt^2a-iCre^* animals, though a small level of aberrant expression was still detected (Extended Data Fig. 1b). This observation motivated us to next consider intersectional approaches, which would allow selective and simultaneous targeting of A1 and C1 cells within the same animal. *Pnmt^2a-iCre^;Dbh^Flp^*double transgenic mice were generated such that C1 and A1 cells are defined by differential recombinase expression (C1: Cre and Flp; A1: Flp)(Fig 1a). Subsequent introduction of AAV-based intersectional tools with distinguishing Boolean logics (CreANDFlp: Con/Fon; FlpNOTCre: Coff/Fon) into RVLM would then allow selective targeting of C1 and A1 neurons^43^. To test the utility of this strategy, we first reassessed the distribution of C1 and A1 neuron populations within RVLM, as previous studies have suggested minimal intermingling of these cell groups^32,33,39^. To do this, a mixture of Con/Fon-eYFP and Coff/Fon-mCherry AAVs was injected into RVLM of *Pnmt^2a-iCre^;Dbh^Flp^* mice and brains were processed for standard immunostaining, dual immunostaining and *in situ* hybridization, or tissue clearing (Fig. 1b). Expression of both AAVs was highly restricted to neurons co-expressing tyrosine hydroxylase (TH) within RVLM, a hallmark of catecholaminergic neurons, with a specificity of 95.0% ± 1.2 for Con/Fon (C1 neurons) and 91.1% ± 2.3 for Coff/Fon (A1 neurons) (Fig. 1c, d). Unlike previous studies, which reported substantial spatial segregation between A1 and C1 populations, we observed Con/Fon- and Coff/Fon-labeled C1 and A1 neurons to occupy largely overlapping domains (Fig. 1e). By performing a linear regression on the proportion of eYFP or mCherry positive cells present on each brain section, we estimated considerable probability (∼20-40%) of finding an A1 neuron in a region enriched with C1 neurons (∼-6.5 mm and ∼-6.9 mm from Bregma), suggesting high intermingling of these two cell types (Fig. 1f). To further confirm the specificity of intersectional AAVs to label C1 neurons, *in situ* hybridization was next performed in combination with immunohistochemistry (ISH/IHC)(Fig. 1g). The analysis of ISH/IHC revealed that the expression of Con/Fon-eYFP colocalized with the expression of *Pnmt* mRNA in 98.4 % ± 0.02 of *Th* mRNA positive cells (Fig. 1h). The extensive spatial overlap of C1 and A1 cells seen in sectioned tissue motivated us to further explore this organization in a three-dimensional setting. To do this, a subset of Con/Fon-eYFP+Coff/Fon-mCherry labeled brains from *Pnmt^2a-iCre^;Dbh^Flp^* mice underwent iDISCO-based tissue clearing, immunolabeling, and registration of labeled A1 and C1 cell bodies (Fig. 1i). Qualitatively, extensive intermingling of C1 and A1 cells was again observed within the intact brainstem, supporting initial observations from sectioned tissue (Fig. 1i-k). Together, these results demonstrate that application of a novel intersectional approach provides selective and simultaneous targeting of A1 and C1 neurons *in vivo*. Moreover, these data suggest that C1 and A1 cells are more intermingled in RVLM than previously reported^33^, highlighting the utility of intersectional approaches to resolve the connectivity and function of these two neuronal populations.

**Figure 1.**
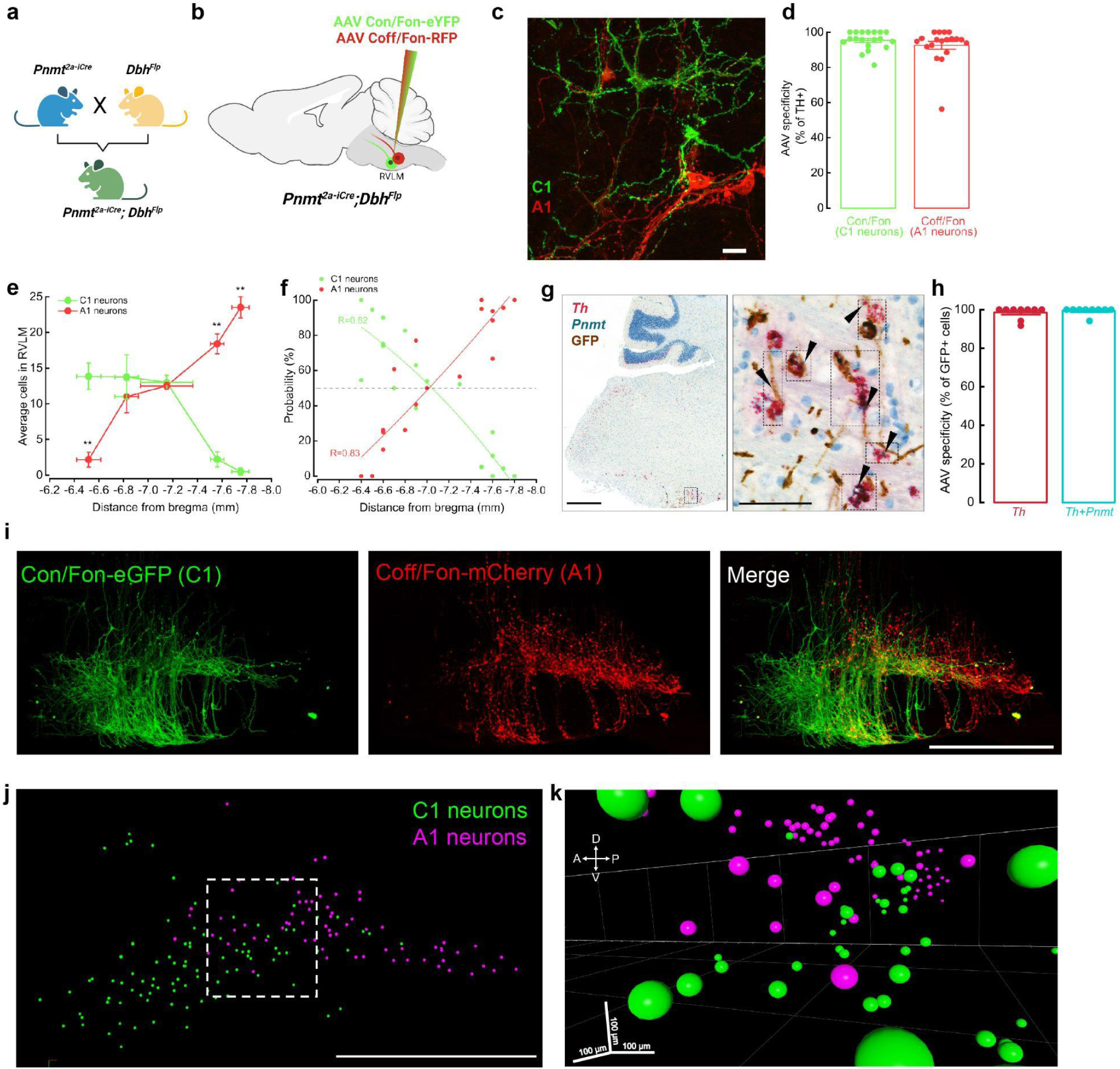
Validation of intersectional approaches to target catecholaminergic C1 neurons. **a,** Breeding strategy for *Pnmt^2a-iCre^*;*Dbh^Flp^* mice. **b,** Schematic of Con/Fon and Coff/Fon viral strategies to label C1 and A1 neurons in RVLM. **c,** Representative image of GFP^+^ cells C1 neurons (green) intermingled with mCherry^+^ A1 neurons (red)(∼-6.8 mm from Bregma). Scale bar: 25 μm. **d,** Average percentage of GFP- or mCherry-labeled cells that co-express TH (n=19 sections from 3 animals). **e,** Average number of GFP- and mCherry-labeled neurons counted throughout RVLM (n=3 animals). Data represents mean ± SEM. *p<0.05, Kruskal Wallis test with Dunn’s post hoc test. **f,** Plot showing the estimated probability of C1 or A1 presence throughout RVLM (n=19 sections from 3 animals). **g,** Representative brightfield images of dual *in situ* hybridization and immunohistochemistry labeling of RVLM. mRNA probes: *Th* (red), *Pnmt* (teal). Antibody: anti-GFP (brown). Scale bars 500 and 100 μm (inset). **h,** Average percentage of GFP-labeled neurons in RVLM co-expressing *Th* or *Th* and *Pnmt* mRNA (n=9 sections from 3 animals). **i,** Representative image of Con/Fon-GFP- and Coff/Fon-mCherry-labeled neurons in RVLM of cleared brain tissue (n=3 animals). **j,** Representative image depicting manual segmentation of C1 and A1 neurons shown in (i). **k,** 3D image from (j) of C1 (green) and A1 (magenta) neuron segmentation highlights intermingling of these cell types in multiple dimensions.

### Diverse synaptic input onto RVLM C1 neurons defined by intersectional transsynaptic rabies tracing

Previous circuit mapping studies have indicated that RVLM C1 and A1 cells receive enriched input from hindbrain areas such as nucleus of the solitary tract (NTS), caudal ventrolateral medulla (CVLM), reticular formation, hypothalamus, ventromedial prefrontal cortex and PAG, in alignment with a proposed role for these cells in modulating physiological responses^33,44^. However, as previous strategies may have lacked synaptic^30,44,45^ or cell-type^33^ specificity, brain-wide input connectivity onto RVLM C1 neurons remains unknown. To address this, we performed intersectional transsynaptic rabies tracing whereby Con/Fon AAVs expressing TVA-mCherry and optimized glycoprotein (oPBG) were expressed in C1 neurons via injection into RVLM of *Pnmt^2a-iCre^;Dbh^Flp^* mice, followed by injection of pseudotyped rabies virus expressing GFP (RABV-GFP) into the same sites 3 weeks later (Fig. 2a). Putative C1 starter cells, or cells that express TVA-mCherry, oPBG, and RABV-GFP, and are thus capable of facilitating transsynaptic spread of rabies virus to presynaptic neurons, were detected exclusively at the injection site (Fig. 2b). In addition, rabies labeling of input neurons was observed throughout the brain in all samples (Fig. 2c, d, Supplementary Video 1). The relative number of input neurons in each brain region, in relation to the total number of rabies-positive cells per whole brain sample, was quantified and classified into three levels: high (cells >10%), moderate (5 %<cells<10 %) and low (cells<5 %). High levels of input neurons were observed in the intermediate reticular nucleus (IRt) and paragigantocellular reticular nucleus (PGi), brain regions involved in respiration, arousal and sleep and pain regulation^46–48^ (Fig. 2c, d). Moderate levels of labeled neurons were found in several areas including the rostral ventrolateral medulla (RVLM), medullary reticular nucleus (RF), pontine reticular nucleus (PnC), gigantocellular reticular nucleus (GRN), and parvocellular reticular nucleus (PCRt), brain regions that participate in locomotion, startle responses, and control of the mandibular and cervical musculature^49–52^ (Fig. 2c, d). Finally, several brain areas displayed low but reliable levels of labeled projections to C1 neurons, including parafacial nucleus (pFRG), periaqueductal gray matter (PAG), inferior olivary complex (ION), superior colliculus (SC), nucleus of the solitary tract (NTS), hypothalamus, and cerebellum (Fig. 2c, d). These regions integrate sensory, somatic (e.g. visual, interoceptive, homeostatic), and motor signals and are involved in modulating respiration, anxiety, and aversion^13,53–56^(Fig. 2c, d). Taken together, these data indicate that C1 neurons receive a complex array of inputs from a wide variety of brain regions, particularly from nuclei involved in the transmission and processing of internal signals and emotional behaviors, suggesting that C1 neurons may function as a brainstem interoceptive hub.

**Figure 2.**
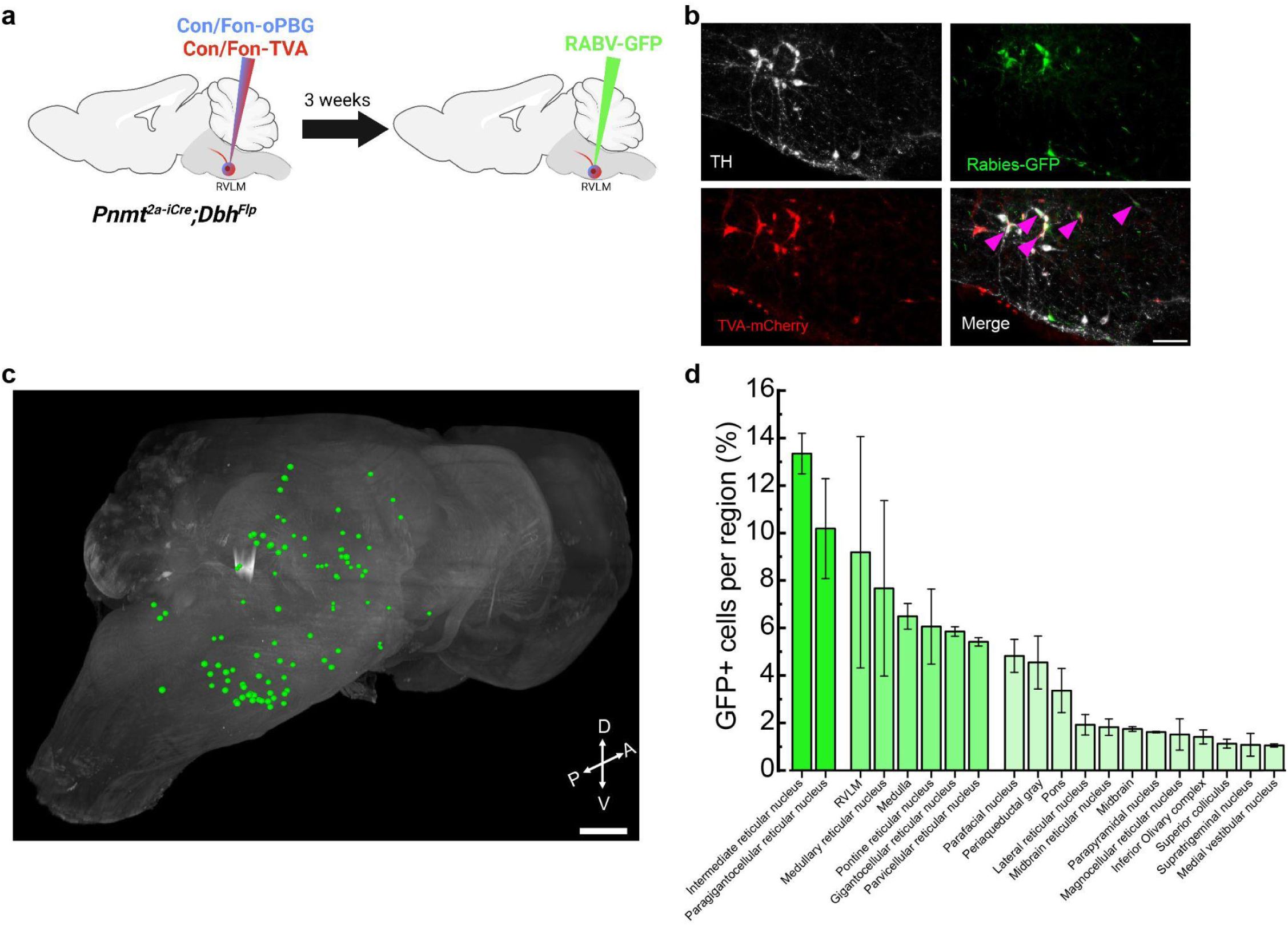
Rabies-mediated transsynaptic tracing identifies presynaptic afferents to C1 neurons. **a,** Strategy for intersectional expression of TVA receptor and optimized glycoprotein (oPBG) in C1 neurons to allow rabies-mediated tracing. **b,** Representative images of C1 starter cells (magenta arrowheads) that are co-labeled by TH (white), GFP (green), and TVA-mCherry (red). Scale bar: 100 μm. **c,** Representative image of a cleared brain showing manual segmentation of rabies-labeled input neurons (green). **d,** Average percentage of rabies-labeled neurons detected in specified brain areas. Data represents mean ± SEM. n=3 animals for b-d.

### Optogenetic stimulation of A1/C1 or C1 neurons enhances anxiety-like behaviors

While research has largely focused on the role of putative C1 neurons over physiological parameters, the consequences of altering C1 neuronal activity on complex behavior remain largely unassessed. Furthermore, much of our understanding regarding the functional role of these cells derives from studies performed in anesthetized animals, a condition that is unlikely to reveal potential involvement in modulating naturalistic behaviors^34^. To address this, we generated experimental cohorts where eYFP-tagged channelrhodopsin (ChR2) was expressed collectively in RLVM A1/C1 cells (Flp-dependent ChR2-eYFP AAV in *Dbh^Flp^* mice) or selectively in C1 cells (Con/Fon-ChR2-eYFP AAV in *Pnmt^2a-iCre^;Dbh^Flp^* mice). Control groups consisted of *Pnmt^2a-iCre^;Dbh^Flp^*mice injected with Con/Fon-eYFP AAV (Fig. 3a). After fiber optic implantation into RVLM and sufficient time for recovery, mice were placed in an open field arena (OF) for testing (Fig. 3a). OF is a commonly used test to measure locomotion, exploration, and anxiety-like behavior in rodents^57,58^. Control and experimental mice were placed into the OF and received optogenetic stimulation at 20 Hz for 30 seconds, followed by a 30 second rest, over a time course of 5 minutes^59^. Control animals spent comparable amounts of time at the corners and in the center region of the OF (Fig. 3b, c). In contrast, bulk activation of RVLM C1 and A1 neurons (A1/C1), as well as specific stimulation of C1 neurons, caused a significant increase in the time that mice spent in the corners of the OF, a phenotype of increased anxiety (Fig. 3b, c). The distance traveled by mice in all groups was similar, with C1-stimulated animals showing a trend towards increased mobility (Fig. 3d). Of note, A1/C1-stimulated mice frequently displayed a transient immobile behavior during light delivery, an effect that was not observed when C1 neurons were selectively targeted (Supplementary Video 2, 3). Consequently, the average velocity of A1/C1 animals during periods of optogenetic stimulation was significantly reduced compared to control or C1-activated mice (Fig. 3e). These data suggest that stimulation of C1 neurons alone is sufficient to increase anxiety-like behaviors in the OFT. Furthermore, the behavioral arrest that was observed by stimulation of A1/C1 neurons in bulk is likely mediated by A1 neurons in RVLM.

**Figure 3.**
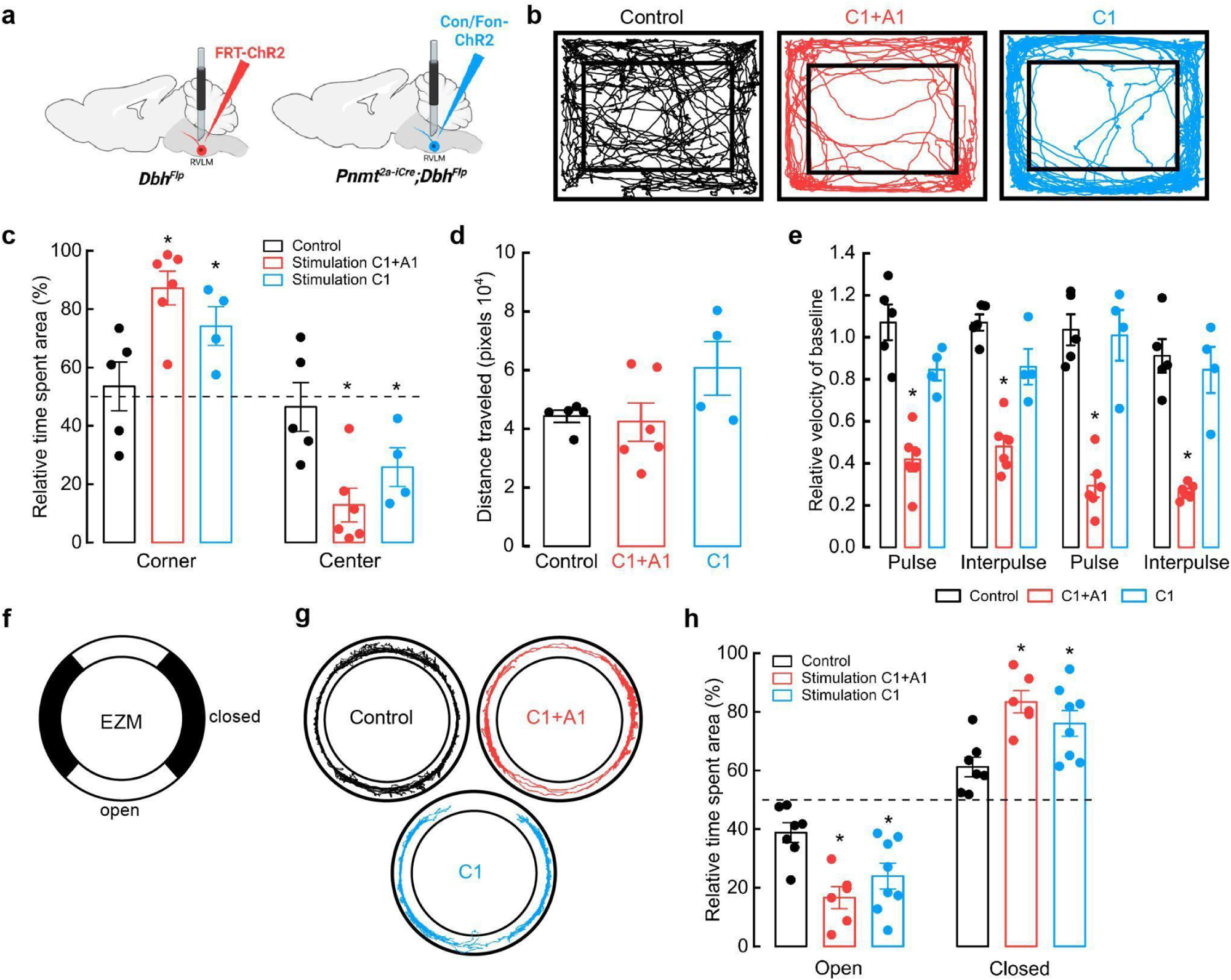
Optogenetic activation of C1 neurons enhances anxiety-like behaviors. **a,** Strategy for targeted viral expression of channelrhodopsin (ChR2) in C1+A1 (red) or C1 (blue) neurons and optic fiber implantation in RVLM. **b,** Representative cumulative trajectories (10 min) in the open field test (OFT) for control (black), C1+A1 (red) and C1 (blue) mice. **c,** Average percentage of time spent in the corners or center of the OFT. **d,** Average total distance traveled in the OFT. **e,** Average change in velocity relative to baseline during (pulse) or between (interpulse) periods of optogenetic stimulation (20 Hz, 20 ms pulse width, for 30 s, 30 s rest). **f,** Schematic of elevated zero maze (EZM). **g,** Representative cumulative trajectories (10 min) in the EZM while optogenetically stimulated (20 Hz, 20 ms pulse width, for 30 s, 30 s rest, 5 min total) for control (black), C1+A1 (red) and C1 (blue) mice. **h,** Average time spent in the open and closed areas of the EZM. Control n=5, C1+A1 n= 6, C1 n=4 animals for b-e; Control n=7, C1+A1 n= 6, C1 n=8 for g, h. Data represents mean ± SEM. *p<0.05 vs control, Kruskal Wallis test with Dunn’s post hoc test.

While the OF is often used to measure anxiety-like behaviors, some have reported behavioral discrepancies with other anxiety-related assays, highlighting the importance of additional testing. The elevated zero maze (EZM), an improved version of the elevated plus maze, is another commonly used assay to assess anxiety-like behaviors in rodents^60^. To ask how selective activation of C1 neurons influences EZM performance, control and experimental mice received optogenetic stimulation (20 Hz, 20 ms, for 30 seconds, 30 seconds rest, for a total of 5 minutes) throughout a 20 min session on the EZM (Fig. 3f). Like OF testing, control mice spent roughly equivalent amounts of time in the open and closed areas of the EZM (Fig. 3g, h). In contrast, activating A1/C1 or C1 neurons caused a significant reduction in the time that mice spent in the open areas, again indicating increased anxiety in these animals (Fig. 3g, h). We did not observe significant differences in the traveled distances and average velocities among all the groups, although A1/C1- and C1-stimulated mice displayed a tendency to travel less and at a slower velocity (Extended Data Fig. 2a, b). These results suggest that stimulation of C1 neurons also increases anxiety-like behaviors in the EZM, confirming our previous findings from OF testing.

### A1 and C1 neurons promote anxiety-like behaviors via a projection to PAG

Our results showing that C1 neurons modulate anxiety-like behaviors motivated us to further explore circuit mechanisms for how this effect might be mediated. Previous studies found that C1 and A1 neurons within RVLM send projections to several brain regions, including PAG. PAG is a key neural structure for the integration of bodily functions, such as cardiovascular and respiratory regulation, as well as emotional behaviors like anxiety and aversion^13,14,19,21,25,29,61,62^. Using intersectional approaches, we first wanted to resolve anatomical characteristics of C1 and A1 axons innervating PAG. A mixture of Con/Fon-eYFP and Flp-dependent-mCherry AAVs were injected into RVLM of *Pnmt^2a-iCre^;Dbh^Flp^* mice to distinguish C1 cells (YFP and mCherry-expressing) from the surrounding A1 cells (mCherry-expressing) (Fig. 4a). Qualitatively, we observed that both C1 and A1 axons innervate PAG, mostly in the ventrolateral (vlPAG) and ventral (vPAG) regions of this nucleus (Fig. 4b). Notably, A1 and C1 axons diverged within PAG, with A1 axons most prominent at the rostral portion of PAG and along the limit of the aqueduct, and C1 axons more concentrated towards the center of lPAG and vlPAG columns, suggesting that C1 and A1 neurons may utilize distinct neural circuit organization motifs to communicate with PAG.

**Figure 4.**
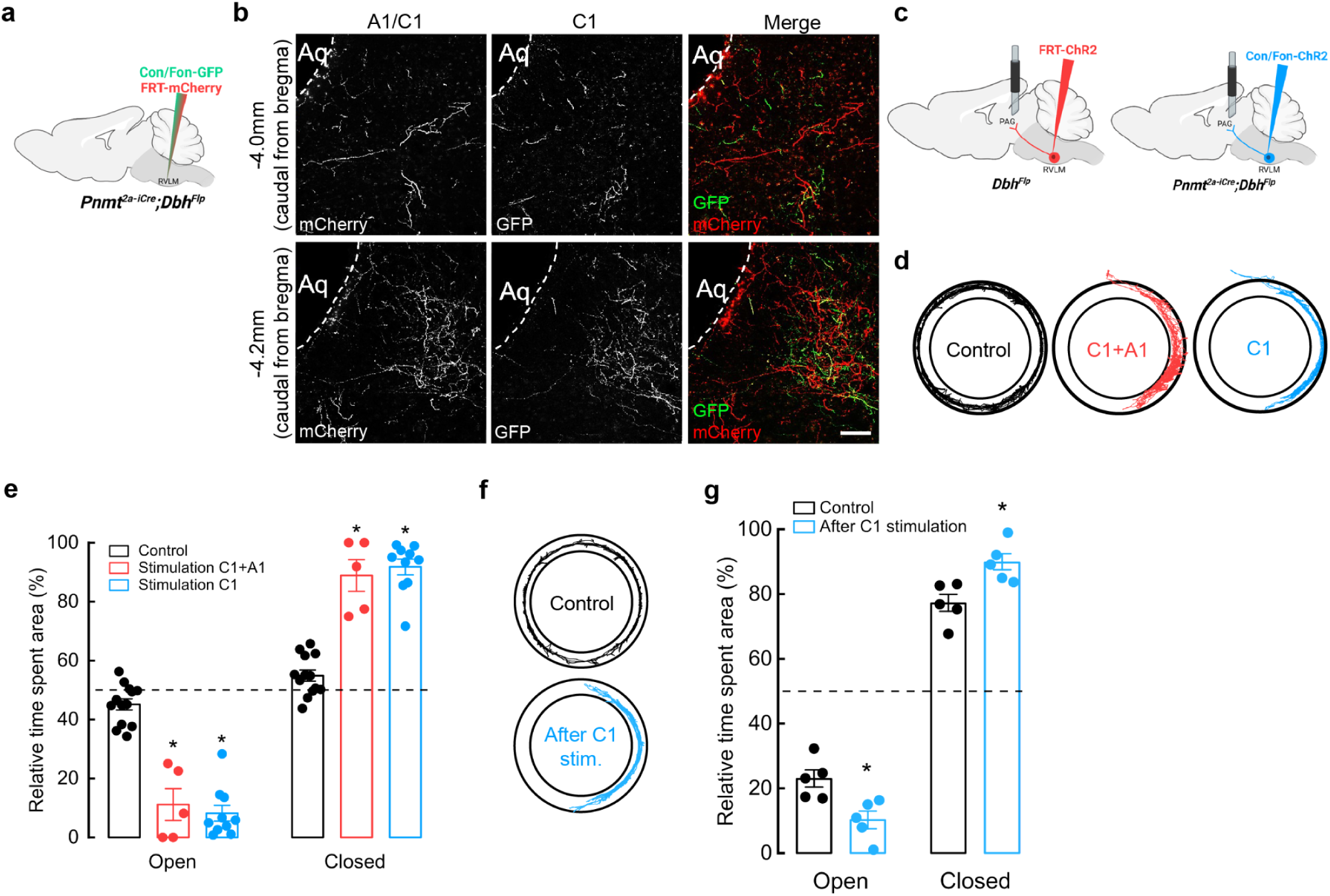
Optogenetic activation of C1 axons innervating vlPAG enhances anxiety-like behaviors. **a,** Schematic of viral strategies to label C1 and A1 neurons in RVLM. **b,** Representative image shows Con/Fon-GFP- and FRT-mCherry-labeled axons in vlPAG (at -4.0 mm and -4.2 mm Bregma). Scale bar: 50 μm. **c,** Strategy for targeted viral expression of channelrhodopsin (ChR2) in C1+A1 (red) or C1 (blue) neurons and optic fiber implantation in PAG. **d,** Representative cumulative trajectories (10 min) in the EZM for control (black), C1+A1 (red) and C1 (blue) mice. **e,** Average percentage of time spent in the open or closed areas of the EZM ((20 Hz, 20 ms pulse width, 30 s stimulation, 30 s rest, 5 min total). **f,** Representative cumulative trajectories (10 min) in the EZM for control (black) or C1 (blue) mice 7 days post optogenetic stimulation at PAG in their homecage (20 Hz, 20 ms pulse width, 30 s stimulation, 30 s rest, 10 min total). **g,** Average percentage of time spent in the open or closed areas of the EZM. n=4 animals for b. Control n=13, C1+A1 n=5, C1 n=10 animals for d, e; Control n=5, C1 n=5 for f, g. Data represents mean ± SEM. p<0.05* vs control, Kruskal Wallis test with Dunn’s post hoc test.

To ask if activation of A1 and C1 projections within PAG is sufficient for promotion of anxiety-like behaviors, we injected a Flp-dependent ChR2-mCherry AAV into RVLM of *Dbh^Flp^* mice (to target A1 and C1 neurons) or a Con/Fon-ChR2 AAV into *Pnmt^2a-iCre^; Dbh^Flp^*mice (to target C1 neurons), followed by optic fiber implantation into vlPAG (Fig. 4c). Additional *Pnmt^2a-iCre^; Dbh^Flp^*mice were injected with AAV Con/Fon-eYFP into RVLM as controls. After recovery, mice were tested in the EZM while A1/C1 or C1 projections within vlPAG (A1/C1^PAG^ or C1^PAG^) were optogenetically stimulated (20 Hz, 20 ms, for 30 seconds, 30 seconds rest, for a total of 5 minutes) during a 20 min session on the EZM. Control mice spent a similar amount of time in the open and closed areas of the maze, while A1/C1^PAG^ and C1^PAG^ mice spent significantly more time in closed areas (Fig. 4d, e). Optogenetic stimulation of A1/C1^PAG^ or C1^PAG^ axons also induced reduction of locomotor activity in all experimental animals (Extended Data Fig. 2c) and reduction in average velocity in C1^PAG^ mice (Extended Data Fig. 2d), indicating that activation of C1 neurons alone is sufficient to promote anxiety-like responses.

Unexpectedly, we noticed that experimental mice displayed signs of elevated stress, such as tail flicking and exacerbated escape behaviors during handling, that persisted many days after completion of optogenetic experiments. This led us to hypothesize that transient activation of C1 neurons might trigger the development of chronic anxiety-like behaviors in these animals. To directly test this, a new set of *Pnmt^2a-iCre^; Dbh^Flp^* mice received injections of either Con/Fon-eYFP AAV (control) or Con/Fon-ChR2 AAV (experimental) into RVLM, to selectively infect C1 neurons, followed by optic fiber implantation into vlPAG. After allowing sufficient time for recovery, both groups received optogenetic stimulation in their home cage (20 Hz, 20 ms, for 30 seconds, 30 seconds rest, for a total of 10 minutes), after which they were immediately returned to the housing rack and left undisturbed for seven days. All mice then underwent EZM testing. Indeed, animals that had received C1^PAG^ optogenetic activation in their home cage one week prior spent significantly less time in the open areas of the EZM compared to control animals (Fig. 4f, g). There were also no significant differences in average velocity or distance traveled between experimental and control groups (Extended Data Fig. 2e, f). These results suggest that heightened activity of C1 projections within vlPAG can facilitate functional adaptations to promote chronic enhancement of anxiety-like phenotypes.

### Optogenetic stimulation of A1/C1^PAG^ and C1^PAG^ axons produce aversion in mice

As previously mentioned, one function of PAG is to promote aversion, an evolutionary mechanism by which perception of harmful stimuli promotes negative emotions, such as anxiety and fear, and subsequent avoidance of such stimuli^19,62–64^. Pathological aversive behaviors are also associated with anxiety disorders^65^. To test whether the activation of A1/C1^PAG^ and C1^PAG^ axons produces aversion, A1/C1^PAG^ and C1^PAG^ mice underwent a real-time place preference assay (RTPP)^66^ where optogenetic stimulation (20 Hz, 20 ms pulse duration) was delivered whenever mice entered a specific side of a two-sided chamber (Fig. 5a, b). Whereas control mice spent roughly equal time on both sides of the chamber, A1/C1^PAG^ and C1^PAG^ mice spent more time in the unstimulated portion of the box, indicating that they found the stimulus to be aversive (Fig. 5b, c). No differences were observed between groups regarding number of entries into the stimulation side (Extended Data Fig. 3a). Additionally, control and C1^PAG^ mice traveled similar distances during the assay, a value that was reduced in A1/C1^PAG^ animals (Fig. 5d). No significant differences were observed in the average velocity between groups (Fig. 5e).

**Figure 5.**
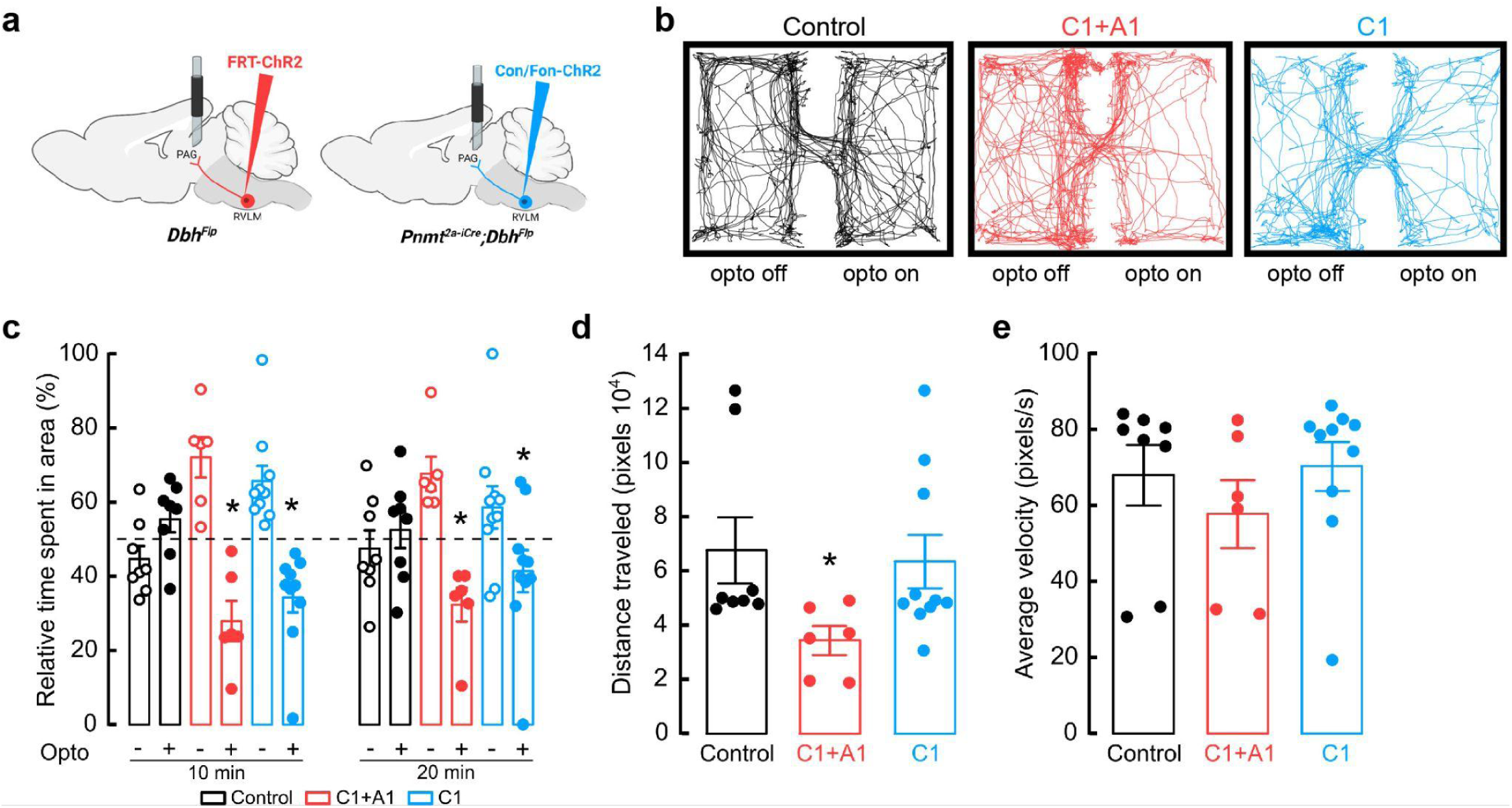
Optogenetic activation of C1 axons innervating vlPAG is aversive. **a,** Strategy for targeted viral expression of channelrhodopsin (ChR2) in C1+A1 (red) or C1 (blue) neurons and optic fiber implantation in PAG. **b,** Representative cumulative trajectories of mice (10 min) in the optogenetic real time place preference (RTPP) assay. Light pulses (20 Hz, 20 ms pulse width) were delivered only when mice inhabited a defined side of the behavioral chamber. **c,** Average percentage of time spent in the unstimulated (opto ‘off’; open circles) or stimulated (opto ‘on’; closed circles) sides of the chamber. **d,** Average distance traveled in RTPP assay. **e,** Average animal velocity during the RTPP assay. Control n=8, C1+A1 n=6, C1 n=10 animals for b-e. Data represents mean ± SEM. *p<0.05 vs control, Kruskal Wallis test with Dunn’s post hoc test.

Because anxiety is associated with changes in respiratory parameters, and our previous results indicate that aversion and anxiety-like behaviors are due to the action of C1^PAG^ axons, we also explored whether stimulation of this connection produced alterations in respiration. Using whole-body plethysmography, we found that optogenetic stimulation of C1 axons innervating PAG increases breathing rate without affecting inspiration or expiration time (Extended Data Fig. 4a-c). Collectively, our behavioral testing show that activation of C1 neurons produces aversion, enhances anxiety-like behaviors, and increases breathing frequency via a direct connection with PAG, demonstrating a novel role for this subcircuit in the promotion and integration of physiological and behavioral anxiety-like symptoms.

### Anxiogenic stimuli activate C1 neurons

A1/C1 neurons respond to a wide variety of stimuli (e.g. hypoxia, hypovolemia, restraint), but are suggested to be less responsive to psychological threats ^23,36,37^. This contrasts with our optogenetic experiments, where activation of these cells drove anxiety-like behavior, motivating us to reassess the responsiveness of C1 cells in our behavioral assays. C1 neurons were again selectively labeled by injecting Con/Fon-eYFP into RVLM of *Pnmt^2a-iCre^;Dbh^Flp^* mice (Extended Data Fig. 5a). After allowing sufficient time for recovery and viral expression to occur, animals were either kept in their home cage (controls), exposed to the EZM test for 20 minutes, or physically restrained for 30 minutes. Mice were sacrificed approximately 60 minutes after assay completion to allow time for expression of cFos, an immediate early gene whose presence is often used as proxy for neuronal activation^67,68^ (Extended Data Fig. 5b, c). Colocalization of cFos and eYFP signal (indicative of C1 cells) was very low in control mice, but robustly increased in response to EZM and restraint (Extended Data Fig. 5b, c). A caveat of these experiments is the static nature and imprecise correlation of cFos labeling with endogenous neural activity. To better visualize the responsiveness of C1 neurons to anxiogenic stimuli in awake-behaving animals, we next expressed Con/Fon-GCaMP8m AAV in C1 neurons of *Pnmt^2a-iCre^;Dbh^Flp^* mice, followed by optic fiber implantation into RVLM for fiber photometry-based bulk calcium imaging (Fig. 6a). Mice were again monitored in the EZM and C1 responses were aligned with behavioral tracking to infer whether alterations in C1 neuronal activity correlated with particular behavioral patterns. We found that C1 neuronal activity sharply increased as mice transitioned from safe (closed) to anxiogenic (open) areas of the EZM, after which calcium signal returned to baseline values (Fig. 6b). Area under the curve (AUC) analysis revealed that calcium activity in C1 neurons increased when mice are present in the open areas of the EZM (26.91 ± 54.35 Z score/s closed vs 899.2 ± 194.25 Z score/s open), again suggesting that these cells are robustly activated by stressful events (Fig. 6b, c). Moreover, it appears their activity is quickly modulated if potential risks are deemed safe, such as after animals enter the open area of the EZM (Fig. 6b).

**Figure 6.**
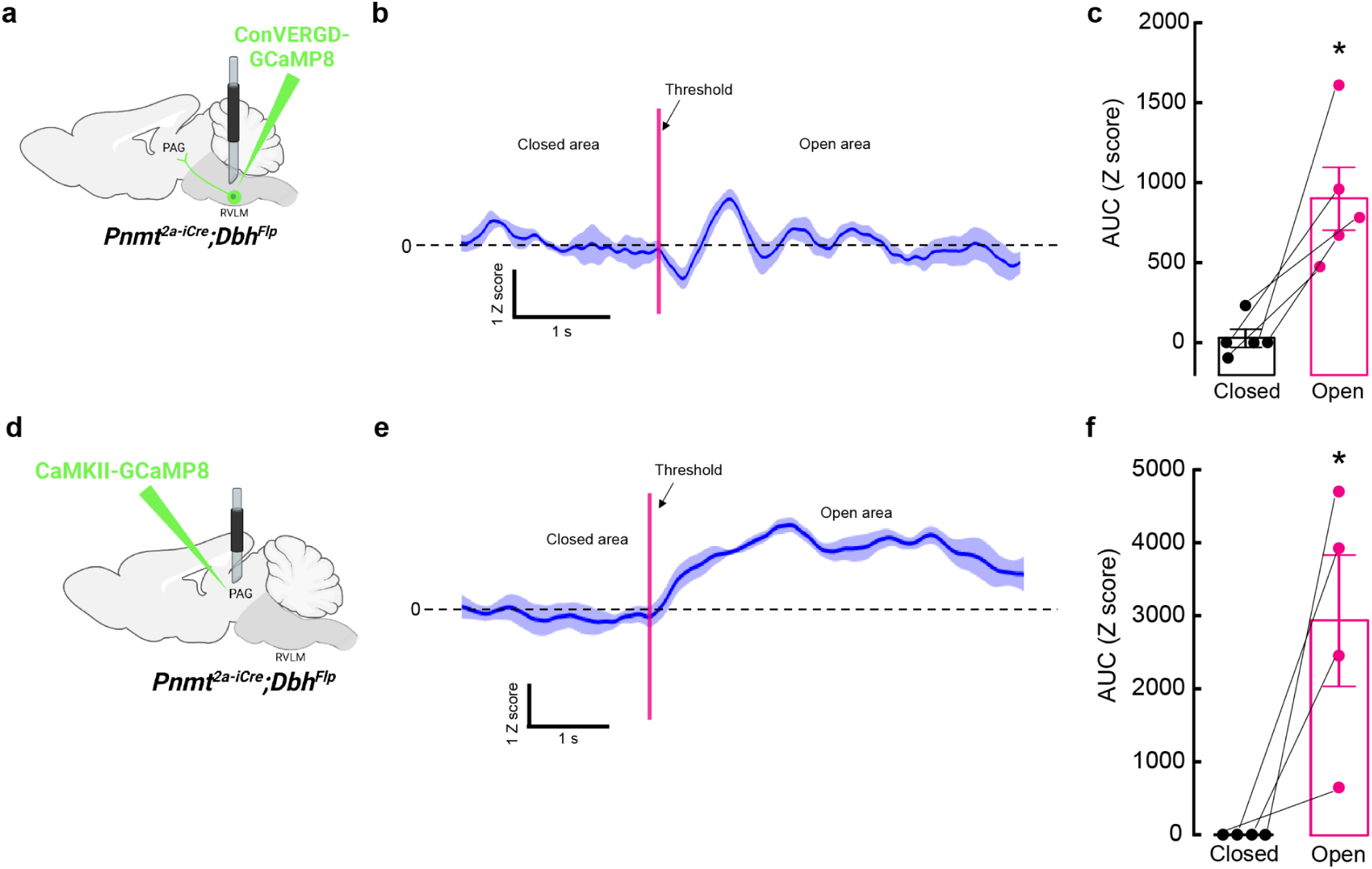
Anxiogenic stimuli activate C1 neurons and excitatory neurons in vlPAG. **a,** Strategy for intersectional viral expression of the calcium indicator GCaMP8m and fiber photometry of C1 neurons. **b,** Representative average calcium activity of C1 neurons while mice transition from the closed to the open area of the EZM. Data represents mean Z score (dark blue) ± SEM (light blue). **c,** Average area under the curve (AUC) analysis of C1 neuron calcium activity from -5 to 0 seconds before crossing the threshold (closed) and from 0 to 5 seconds after entering the open area (open). Data represents mean AUC ± SEM. n=5 animals, *p<0.05, paired t-Test. **d,** Strategy for viral expression of GCaMP8m and fiber photometry from excitatory neurons in vlPAG. **e,** Representative average calcium activity of excitatory vlPAG neurons while mice transition from the closed to the open area of the EZM. Data represents mean Z score (dark blue) ± SEM (light blue). **f,** Average AUC analysis of excitatory vlPAG neurons calcium activity from -5 to 0 seconds before crossing the threshold (closed) and from 0 to 5 seconds after entering the open area (open). Data represents mean AUC ± SEM. n=4 animals, *p<0.05, paired t-Test.

### Excitatory neurons in vlPAG are activated by anxiogenic stimuli

Using intersectional strategies, our experiments found that optogenetic activation of C1 axons in PAG induces aversion and enhances anxiety-like behaviors (Figs. 4, 5). Moreover, prior research suggests that vlPAG neurons process threat information and determine the appropriate behavioral response to such threat^14^. Maladaptation to this function could thus lead to potential threat overestimation and facilitation of pathological anxiety^69,70^. To address how increased C1 activity might influence downstream brain areas, specifically vlPAG, we again performed fiber photometry-based calcium imaging in mice undergoing EZM testing, but now recording from excitatory neurons within vlPAG via expression of CaMKII-GCaMP8m AAV and subsequent fiber implantation into the region (Fig. 6d). Similar to C1 neurons, we observed that basal activity of excitatory vlPAG neurons was low and bound to exploratory movements, with signal increasing when mice transitioned from closed to open areas of the EZM (Fig. 6e). Of note, the anxiogenic-induced activation of excitatory vlPAG neurons was more stable than what was observed for C1 neurons, with increased GCaMP signal lasting for more than 5 seconds post-activation (Fig. 6e). AUC analysis again revealed that excitatory vlPAG neurons are more active in the open areas of EZM, suggesting that they may be selectively recruited in times of enhanced stress, similar to C1 neurons (Fig. 6f).

To address whether C1 neurons influence vlPAG activity, *Pnmt^2a-iCre^;Dbh^Flp^*mice received injections of Con/Fon-ChRmine AAV into RVLM and CaMKII-GCaMP8m AAV into vlPAG. Optic fibers were implanted at both injection sites, allowing simultaneous optogenetic activation of C1 neurons and calcium imaging of excitatory vlPAG neurons (Extended Data Fig. 6a). Optogenetic activation of C1 neurons caused robust calcium responses in excitatory vlPAG neurons (Extended Data Fig. 6b). To verify these findings in awake-behaving animals, we instead expressed Con/Fon-DREADD-Gq-mCherry AAV in C1 neurons (for chemogenetic-based activation) and CaMKII-GCaMP8m AAV in the vlPAG of *Pnmt^2a-iCre^;Dbh^Flp^* mice, along with optic fiber implantation into vlPAG (Extended Data Fig. 6c). Fiber photometry was performed during EZM, before and after the administration of deschloroclozapine (DCZ), a potent and selective DREADD agonist, to transiently activate C1 neurons^71^ (Extended Data Fig. 6d). Without DCZ, basal activity of excitatory vlPAG neurons was low but increased upon transition into the open area of the maze, as in previous experiments (Extended Data Fig. 6d, e). Upon administration of DCZ and return to the EZM, we observed a significant increase in the magnitude of excitatory vlPAG neuronal activity specifically when mice entered the open areas of the maze (Extended Data Fig. 6d, e). Collectively, these data show that activation of C1 neurons facilitates the enhancement of excitatory vlPAG responses to anxiogenic stimuli.

### Chemogenic inhibition of C1 neurons is anxiolytic

Based on our findings that C1 neurons increase their activity under stressful conditions (Fig. 6, Extended Data Fig. 5) and that optogenetic activation of these neurons promotes anxiogenic behavior (Fig. 4, Fig. 5), we hypothesized that selective inhibition of C1 neuronal activity could be advantageous for reduction of stress-related behaviors. However, a challenge with implementing such a strategy is that C1 cell bodies are sparsely distributed throughout RVLM, introducing concerns that light delivery would be insufficient for effective inhibition. We instead utilized inhibitory chemogenetic tools, injecting Con/Fon-hM4D-Gi-mCherry^43^ or Con/Fon-eYFP AAV (for control animals) into RVLM of *Pnmt^2a-iCre^;Dbh^Flp^*mice (Fig. 7a). After allowing sufficient time for viral expression, control and experimental mice were injected with the chemogenetic ligand DCZ 15 minutes prior to testing on the EZM. Control and experimental mice spent similar amounts of time in the open and closed areas of the maze, implying that reduction of C1 activity did not significantly alter EZM-related behavior in this setting (Fig. 7b, c). Traveled distance and average velocity also remained unaltered upon inhibition of C1 neurons (Extended Data Fig. 2g, h).

**Figure 7.**
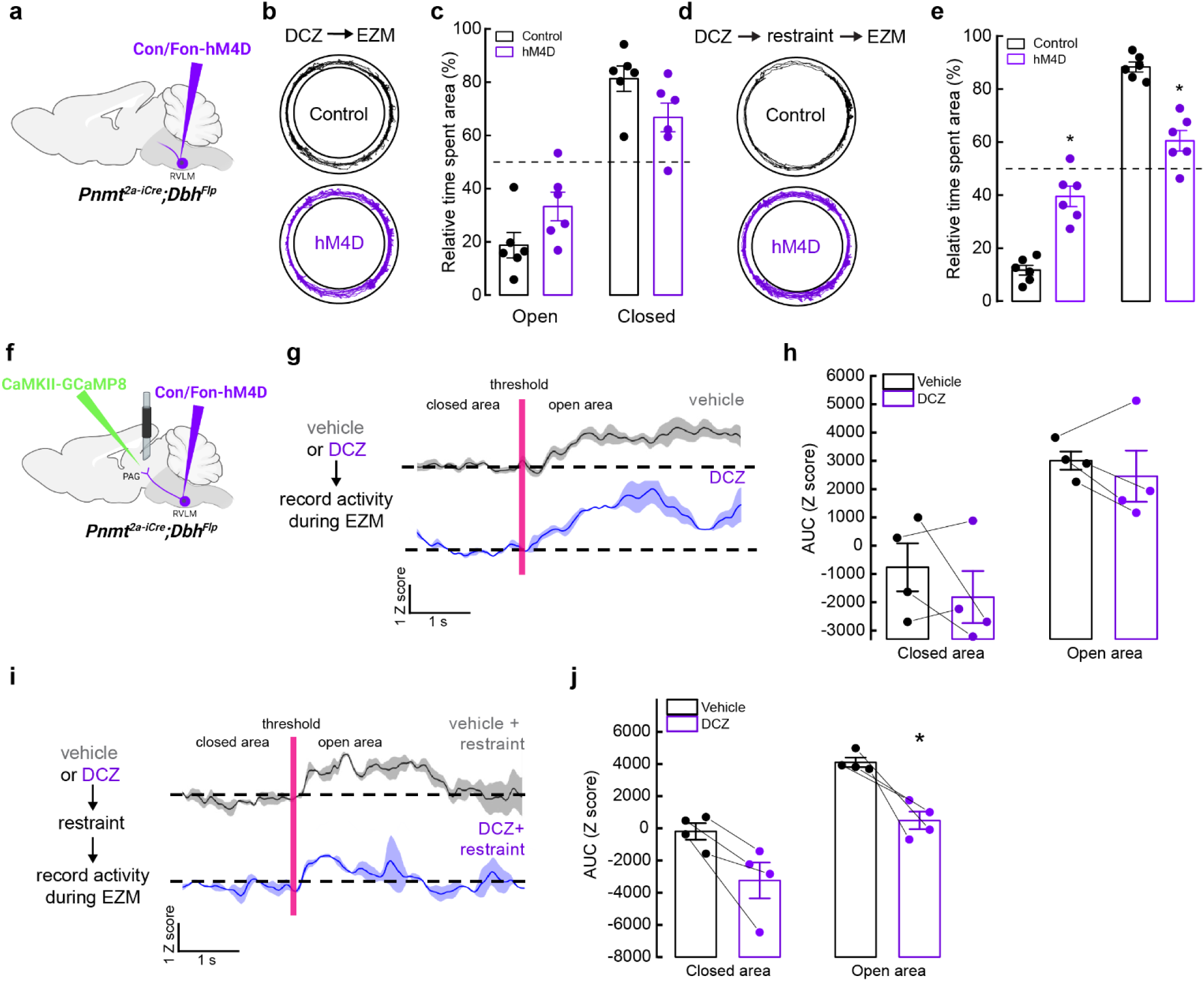
Inhibition of C1 neurons is anxiolytic and reduces vlPAG activity in response to stress. **a,** Strategy for intersectional expression of the inhibitory chemogenetic receptor hM4D in C1 neurons. **b,** Representative cumulative trajectories (10 min) in the EZM for control (black) and hM4D (purple) mice after administration of the DREADD agonist DCZ. **c,** Average percentage of time spent in the open or closed areas of the EZM. **d,** Representative cumulative trajectories (10 min) in the EZM for control (black) and hM4D (purple) mice after administration of the DREADD agonist DCZ and restraint (30 min). **e,** Average percentage of time spent in the open or closed areas of the EZM following restraint. **f,** Strategy for viral expression of GCaMP8m and fiber photometry from excitatory neurons in vlPAG and intersectional expression of hM4D in C1 neurons. **g,** Representative average calcium activity of excitatory vlPAG neurons while mice transition from the closed to the open area of the EZM, in mice pretreated with vehicle (grey) or DCZ (purple). Data represents the mean (dark line) ± SEM (light line). **h,** Average AUC analysis of excitatory vlPAG neuron calcium activity from -5 to 0 seconds before crossing the threshold (closed) and from 0 to 5 seconds after entering the open area (open). Data represents mean ± SEM. **i,** Representative average calcium activity of excitatory vlPAG neurons while mice transition from the closed to the open area of the EZM, in mice pretreated with vehicle (grey) or DCZ (purple) before undergoing restraint (30 min) and EZM. Data represents the mean Z score (dark line) ± SEM (light line). **j,** Average AUC analysis of excitatory vlPAG neuron calcium activity from -5 to 0 seconds before crossing the threshold (closed) and from 0 to 5 seconds after entering the open area (open). Data represents mean AUC ± SEM. Control n=6, hM4D n=6 for b, c; Control+restraint n=6, hM4D+restraint n=6 for d, e; Vehicle n=4, DCZ n=4 for g-j. *p<0.05, Kruskal Wallis test with Dunn’s post hoc test.

While selective inhibition of C1 neurons did not alter immediate behaviors, we wondered if targeting these cells could still reduce the promotion or severity of anxiety-like behaviors over longer timescales, as neuromodulators and neuropeptides can have slower dynamics of action^72^. Thus, we implemented a behavioral paradigm where mice were restrained for 30 minutes prior to placement in the EZM. This action was previously shown to exacerbate stress-related behaviors in EZM, a phenotype that was confirmed in our animals^73^ (Extended Data Fig. 7a, b). Next, Con/Fon-hM4D-Gi-mCherry (experimental) and Con/Fon-eYFP (control) AAV mice received DCZ prior to undergoing 30 minutes of physical restraint, after which they were immediately tested on the EZM. As anticipated, control mice spent almost all their time in the closed portions of the maze, indicating exacerbated anxiety due to prior restraint exposure (Fig. 7d, e). Strikingly, this effect was significantly mitigated by inhibition of C1 neurons during restraint, with hM4D-expressing mice performing similarly in EZM to what we previously observed in unrestrained animals (Fig. 7d, e). C1-inhibited mice also increased their speed and distance of travel compared to control animals (Extended Data Fig. 7i, j).

We then asked whether inhibition of C1 neurons could also alter activity in vlPAG. Con/Fon-hM4D-Gi-mCherry AAV was injected into RVLM and CaMKII-GCaMP8m was injected into vlPAG of *Pnmt^2a-iCre^;Dbh^Flp^* mice, along with implantation of an optic fiber in vlPAG for fiber photometry-based calcium imaging (Fig. 7f). Excitatory vlPAG responses to the open area of the EZM were first collected with administration of saline (vehicle) or DCZ, with no significant difference observed between conditions (Fig. 7g, h). Next, we repeated our previous strategy where mice received saline or DCZ prior to undergoing restraint for 30 minutes. In line with previous findings, we observed that inhibition of C1 neurons during restraint reduced the responsiveness of excitatory vlPAG neurons to the anxiogenic stimuli of the EZM (Fig. 7i, j). Taken together, these data suggest that C1 neurons contribute to the promotion of heightened states of anxiety via activation of excitatory vlPAG neurons, and that disruption of this signaling can be anxiolytic for future stressful events.

## Discussion

Previous studies indicate that catecholaminergic neurons located within RVLM, specifically A1 and C1 cells, promote a wide range of physiological functions, including blood pressure regulation, breathing, and glycemia modulation. Here, we report that C1 neurons also potently modulate cognitive processes, specifically anxiety-like behavior, and we identify an underlying circuit by which this effect occurs. These experiments necessitated the development of a new transgenic mouse line (*Pnmt^2a-iCre^*) in combination with novel intersectional strategies to unambiguously target C1 neurons. Transsynaptic rabies tracing from C1 neurons indicates that they receive direct input from a variety of brain regions across motor, visceral, sensory, and somatic domains, many of which were previously undefined (Fig. 2). Activation of C1 neurons also potentiates anxiety-like behaviors (Fig. 3), an effect that was replicated by selective stimulation of C1 axons in vlPAG (Fig. 4), implicating this subcircuit (C1→vlPAG) as a driver of psychological stress for the first time. Activation of C1 cells is aversive to mice, as seen by their avoidance of areas where ChR2-expressing C1 cells were photostimulated (Fig. 5). Notably, chemogenic inhibition of C1 cells did not alter EZM-related performance in real time but was instead effective at mitigating the enhancement of anxiety following highly stressful events, suggesting that this circuit could be therapeutically relevant for alleviation of anxiety-like symptoms.

### Intersectional strategies facilitate access to catecholaminergic subtypes in RVLM

RVLM contains the largest cluster of adrenergic neurons in the brain, the C1 cells, which are in close proximity to another catecholaminergic subtype, the norepinephrine-expressing A1 cells^31,40,74^. Previous studies have suggested that A1 and C1 intermingling is minimal and largely restricted to the boundary of rostral and caudal VLM, known as the intermediate VLM (iVLM)^33,40^. To achieve specificity for targeting C1 cells *in vivo*, researchers have thus relied on anatomical delineation and molecular tools targeted to catecholaminergic cells, such as the artificial *Phox2b*-specific promoter PRSx8^32,36,59,75,76^, Cre-expressing transgenic rodent lines (*Th^Cre^* and *Dbh^Cre^*)^33,35,37,39,59,77^, or saporin fused to DBH antibody^78,79^. Using these types of approaches, C1 neurons have been attributed to the promotion of a wide range of physiological processes, such as arousal, breathing, inflammation, glycemia, and blood pressure regulation^32,35,37,39^. However, an issue with these tools is that while intended to selectively target catecholaminergic neurons, they can lack cell type specificity^40,76^, and critically, do not distinguish between A1 and C1 cells^35,36,38,59^. It is therefore difficult to ascertain from previous studies whether roles ascribed to C1 cells are accurate or involve nearby A1 cells for promotion. Here, application of novel strategies (*Pnmt^2a-iCre^*;*Dbh^Flp^*mice in combination with intersectional AAV tools) allowed us to unambiguously target C1 neurons within the rodent RVLM for the first time, an important advance as A1 and C1 neurons appear to be more intermingled than previously suggested (Fig. 1).

### Brainwide C1 circuit organization revealed by transsynaptic rabies tracing

Leveraging our intersectional model to perform transsynaptic rabies tracing, we observed input labeling in several brain regions that align with previous findings obtained using traditional strategies, including IRt, PGi, MRt, NRG, PnC, PCR, RTN, RVLM and PAG^33,80–84^ (Fig. 2). The ability of rabies tracing to replicate known connectivity onto RVLM catecholaminergic neurons not only supports the accuracy of this tool, but also sets the stage for future experiments to functionally characterize these connections using next-generation rabies-based reagents. Specifically, while the B19 strain of rabies used for this study is typically limited to anatomical-based labeling, due to the rapid toxicity of the virus, intersectional rabies tracing with the less toxic N2c strain was also recently developed^85^. Thus, it should be feasible to apply N2c-based rabies tracing to C1 cells in the future, using strains that express molecules to monitor (via GCaMP8) or alter (via excitatory or inhibitory opsins) pre-synaptic inputs, to uncover exactly how defined inputs influence C1 activity^86^. It is also important to note that while our rabies tracing corroborates many previously defined afferents to C1 cells, the cell type identity of these inputs remains largely undefined. However, the reduced toxicity of the N2c strain should allow straightforward transcriptional profiling of rabies-labeled inputs, an important future step towards understanding the functional organization of C1 neural circuits.

The increased efficiency of rabies tracing over traditional molecular tools also facilitated discovery of several novel inputs onto C1 cells, notably the lateral parafacial region (pFRG) and the pedunculopontine nucleus (PPN). The pFRG consists of a group of glutamatergic neurons within the RTN nucleus, located ventrolateral of the facial nucleus. These neurons control breathing by integrating signals from other brain regions, particularly during physical stress^87,88^. The PPN is located in the upper brainstem and is composed of at least three cell types; glutamatergic, cholinergic, and GABAergic. This brain region is involved in several functions including arousal and motivation, and also thought to facilitate motor-related deficits in Parkinson’s disease^89–91^. Future studies are necessary to interrogate the roles that each of these regions (and the cell types within) have in modulating C1 activity but should be attainable by building on the intersectional strategies reported here.

### C1 neurons promote anxiety-like behaviors

The breadth of studies characterizing C1 function are almost exclusively relegated to the measurement of physiological responses. Here, using new methods that unequivocally target C1 cells for the first time, we re-evaluated the functional impact of these neurons, including their involvement in the promotion of anxiety-like behaviors. We observed behavioral changes in mice indicative of increased anxiety, occurring both when A1 and C1 cells within RVLM were optogenetically activated, as well as when C1 cells alone were targeted (Fig. 3). Importantly, combined activation of A1 and C1 cells also produced a transient behavioral arrest in experimental animals (Supplementary Video 2). This phenotype was never observed when C1 cells were selectively activated (Supplementary Video 3), highlighting the utility of our intersectional approach for precise dissection of C1 neural circuit function. We speculate that this transient freezing indicates distinguishable features between RLVM A1 and C1 cells, with the former potentially promoting physiological alterations, such as alterations in blood pressure or respiration, that could facilitate temporary behavioral arrest. Further work is needed to confirm these hypotheses but will require significant expansion of current NOT logic intersectional tools so that A1 cells can be selectively and robustly targeted.

### C1→vlPAG subcircuit is sufficient to induce anxiety

Having established that stimulation of C1 neurons enhances anxiety-like behaviors, we next searched for downstream brain region(s) through which this effect might be mediated. Previous studies have suggested that C1 neurons innervate several brain nuclei involved in promotion of anxiety-like behaviors, including Locus Coeruleus (LC)^73,75,92^, lateral parafacial region (pFRG)^93^ and PAG^13,14,19^. For this study, we chose to focus on PAG due to its multifaceted role processing physiological and psychological components of the stress response, as well as its involvement in anxiety and panic disorders. PAG is subdivided into columns, where each column participates in distinct responses. For instance, it is thought that dorsal PAG (dPAG) facilitates pain modulation and escape responses^28,94^, while lateral and ventrolateral PAG (lPAG and vlPAG) are preferentially involved in freezing and predator avoidance^95^. Here, we show that C1 and A1 neurons predominantly innervate lPAG and vlPAG regions, though the spatial distribution of their terminals within these regions varies (Fig. 4b). Additionally, optogenetic stimulation of C1 or A1/C1 axons in vlPAG (C1^PAG^ or A1/C1^PAG^) is sufficient to enhance anxiety-like behavior similarly to activation of A1 and C1 cell bodies (Fig. 3 and Fig. 4). Of note, stimulation of A1/C1^PAG^ or C1^PAG^ axons did not induce the behavioral arrest that was observed during optogenetic activation of A1/C1 cell bodies, indicating that this phenotype likely depends on recruitment of A1 neurons projecting to brain regions other than PAG.

According to the predatory imminence continuum theory, the type of defensive behavior is determined by how imminent the danger is^96^. Previous work has suggested that vlPAG neurons compute the time and degree of the threat, estimating the probability of the threat and determining the appropriate behavioral response. This function also highlights the importance of this structure in promotion of anxiety disorders, where threat overestimation is a hallmark^14,70^. This led us to speculate that stimulation of C1^PAG^ axons may increase the activity of vlPAG neurons to exacerbate threat assessment and enhance anxiety-like phenotypes. Our fiber photometery-based recordings of excitatory responses in vlPAG when C1 activity is altered support this hypothesis (Fig. 7). However, it must be noted that PAG is composed of molecularly diverse cell types that release a variety of molecules implicated in promotion of aversive phenotypes, such as dynorphin, substance P, enkephalins, and cholecystokinin^95,97^, emphasizing the utility of future studies to clarify downstream targets of C1 projections in vlPAG and their function. Additionally, while our data demonstrate that the C1→vlPAG circuitry is an important driver of anxiety-like behaviors and aversion, we also cannot rule out potential contributions from other brain regions. For instance, activation of the A1/C1 projections to LC (A1/C1→LC) induces alertness and produces arousal^39^ and chronic stress can sensitize LC to subsequent stressors facilitating anxiety-like behaviors^73^. Activation of pFRG inputs to A1/C1 neurons also stimulates active expiration^36,87^, disrupts normal breathing cycles, and potentially contributes to the development of anxiety^93,98^. Undoubtedly, more work is necessary to elucidate the roles of other brain regions up- and downstream of C1 neurons in promotion of aversion and anxiety.

### Inhibition of C1 neurons alleviates severity of subsequent anxiety-like responses

Our observation that activation of C1 neurons enhances anxiety-like behaviors warranted us to explore whether inhibition of these cells would have the opposite effect. We found that it did not, at least not when behavioral testing immediately followed induction of C1 inhibition (Fig. 7b, c). Instead, we observed that reducing the activity of C1 cells during highly stressful events (e.g., restraint stress) significantly reduced the severity of anxiety-like behaviors in subsequent testing (Fig. 7d, e). A similar phenomenon has been reported for noradrenergic neurons in LC, where inhibition of these cells during stressful events averts the development of anxiety-like behaviors^73^. Our results might not be surprising in light of classic and recent studies showing that catecholamines can act over multiple timescales^99,100^. Additionally, functional connectivity between vlPAG and cortical centers seems to depend on stress levels: at rest vlPAG is functionally correlated to fronto-limbic areas but “disconnects” from those structures and increases connectivity with the insula during stress^61^.

Ultimately, our findings indicate that the C1→vlPAG subcircuit is a critical and novel contributor for the enhancement and chronification of anxiety-like behaviors. In addition, inhibition of C1 neurons can be anxiolytic without deleterious side effects, making it plausible to leverage C1-related neural circuits as therapeutic targets for the alleviation of anxiety-related symptoms. Although our findings are limited to mice, functional connections between RVLM and PAG have been observed in humans^101^. Our intersectional approaches enable future studies aimed at understanding the functional impact of C1→PAG connectivity and the contribution of diverse afferents (e.g. interoceptive inputs) on the activity of C1 neurons, important information for understanding how integration of cognitive and physiological information influences the promotion of anxiety disorders.

## Supplementary Video 1

3D reconstruction of a cleared brain from a *Pnmt^2a-iCre^;Dbh^Flp^* mouse previously injected with AAV Con/Fon-oPBG and Con/Fon-TVA-mCherry AAVs and pseudotyped rabies virus expressing GFP (RABV-GFP). Segmented somas from afferent neurons projecting onto C1 neurons are visible as green spheres (manual segmentation).

## Supplementary Video 2

A *Dbh^Flp^* mouse, previously injected with AAV FRT-Chr2-mCherry and implanted with an optic fiber in RVLM, freely moving in the open field (OF) arena while tethered to an optic fiber patch cord. Optogenetic activation of C1 + A1 neurons in RVLM (30-s train at 20 Hz, 20-ms pulse width) temporary causes behavioral arrest.

## Supplementary Video 2

A *Pnmt^2a-iCre^;Dbh^Flp^* mouse, previously injected with AAV Con/Fon-ChR2-eYFP and implanted with an optic fiber in RVLM, freely moving in the open field (OF) arena while tethered to an optic fiber patch cord. Optogenetic activation of C1 neurons in RVLM (30-s train at 20 Hz, 20-ms pulse width) fails to induce behavioral arrest.

## Materials and Methods

### Animals

All animal procedures were approved by the Animal Care and Use Committee of St Jude Children’s Research Hospital. *Pnmt^Cre^* mice were kindly gifted by Dr. Steven Ebert (University of Central Florida) and described previously^42^. *Dbh^Flp^* mice, originally developed in the laboratory of Dr. Patricia Jensen^102^ (National Institute of Environmental Health Sciences), were obtained from Jackson Laboratories (Strain # 033952). *Pnmt^2a-iCre^*mice were generated using CRISPR-Cas9 technology. Briefly, a T2A-iCre sequence (1119 bp) was inserted in place of the stop codon in *mPnmt* exon 3. sgRNAs targeting within 20 bp of the desired integration site were designed with at least 3 bp of mismatch between the target site and any other site in the genome. Targeted integration was confirmed *in vitro* prior to moving forward with embryo injections. C57BL/6J fertilized zygotes were micro-injected in the pronucleus with a mixture of Cas9 protein at 30-60 ng/μL, and single guide RNA at 10-20 ng/μL each, and a ssDNA at 5-10 ng/μL (a ribonucleoprotein complex). The injected zygotes, after culture in M16 or alternatively Advanced-KSOM media, were transferred into the oviducts of pseudo-pregnant CD-1 females. Founder mice were genotyped by targeted next generation sequencing followed by analysis using CRIS.py. *Pnmt^2a-iCre^*;*Dbh^Flp^* double transgenic mice were generated by crossing *Dbh^Flp^* with *Pnmt^2a-iCre^*mice. C57BL/6J mice (∼12 weeks old) were purchased from Jackson Laboratories. Mice were housed at 21-22 °C with 12 h light/12 h dark cycles and had free access to fresh water and standard chow pellets. Adult (6-20 weeks old) male and female mice were used for experiments.

### Stereotaxic Procedures

For all stereotaxic procedures, mice were anesthetized with isoflurane: 2.0 % at 1 L/min for induction; 1-2 % at 250 mL/min for maintenance using a low-flow electronic vaporizer (SomnoFlo, Kent Scientific. Viruses were delivered using stereotaxic equipment (Kopf), a UMP3 microinjection syringe pump (World Precision Instruments) and a pulled glass micropipette. All injections were dispensed at a rate of 50 nL/min and allowed to diffuse for 5 minutes before the micropipette was withdrawn. A 2:1 mixture of virus and mannitol was used to enhance expression and viral spread^103^. To reach the full extent of RVLM, three injections (500 nL each) were performed at three different locations. ML and DV coordinates remain unchanged at 1.4 mm and 4.8 mm from the surface of the brain, respectively. AP coordinates for each location were 2.3 mm, 2.6 mm and 3.16 mm posterior from Lambda. Ventrolateral PAG (vlPAG) was targeted with a single injection (100 nL), using the following coordinates: ML -1.04 mm, 4.1 mm posterior from Bregma and DV 2.55 mm at 14° angle from the surface of the brain. Optic fiber cannulas for optogenetics or fiber photometry were fixed to the skull using superglue (Loctite) and dental cement (Ortho-Jet). Accuracy of virus injections for all samples was confirmed via histology.

### Labeling A1 and C1 neurons

To label A1 and C1 neurons, a 1:1 mixture of *AAV8 Ef1a-CONFON-IC++-eYFP* (Addgene #137155) and *AAV8 Ef1a-CoffFon2-ChR2-mCherry* (Addgene #137134) was injected into the left RVLM of *Dbh^Flp^*; *Pnmt^2a-iCre^* mice. Mice were euthanized and their brains harvested 3-4 weeks after the surgery for subsequent histology.

### Retrograde trans-synaptic tracing

For monosynaptic tracing of inputs to C1 neurons, three stereotaxic injections (500 nL each) of a 1:1 mixture of *AAV8 Ef1a-CONFON2-oPBG*^43^ (Addgene #131778) and *AAV8 Ef1a-CONFON2-TVA-mCherry*^43^ (Addgene #137132) were performed into the left RVLM of *Pnmt^2a-iCre^*;*Dbh^Flp^* mice. For monosynaptic tracing of inputs to A1+C1 neurons, the same procedure was followed using *Dbh^Flp^*mice and a 1:1 mixture of *AAV8 CAG-FLEx-FRT-RG* (St. Jude) and *AAV8 CAG-FLEx-FRT-TC* (St. Jude). Three weeks later, mice received three injections (500 nL each) of *SADΔG-GFP-EnvA* into the same coordinates. Animals were allowed to recover for at least one hour and then were housed individually for 5-7 days before sacrificing.

### Optogenetic stimulation of A1+C1 and C1 neurons

Viral constructs were injected into RVLM as detailed above. To target A1+C1 neurons, *Dbh^Flp^* mice were injected with *AAV1 pAAV-CAG-FLEXFRT-ChR2(H134R)-mCherry* (Addgene #75470). To specifically target C1 neurons, *Pnmt^2a-iCre^*;*Dbh^Flp^*mice were injected with *AAV8 Ef1a-CONFON2-ChR2-mCherry*^43^(Addgene #137132). To stimulate cell bodies or axons projecting to vlPAG, a borosilicate optic fiber cannula (Doric Lenses, N.A. 0.66, 400 μm core) was implanted above the injection site either in RVLM (ML 1.4 mm, AP 2.6 mm posterior to bregma, DV 4.8 mm), or vlPAG (ML -1.04 mm, 4.1 mm posterior from Lambda and DV 2.45 mm at 14° angle). Animals were allowed to recover for 3-4 weeks before experimentation.

### Fiber photometry of C1 neurons

To record neuronal activity of C1 neurons, *Pnmt^2a-iCre^*;*Dbh^Flp^*mice were injected with *AAV8 hSyn-ConVERGD-GCaMP8m-CW3SL*^85^ (St. Jude) into the left RVLM. A fiber optic cannula (Doric Lenses, N.A. 0.66, 400 μm core) was implanted above the injection site in RVLM (ML 1.4 mm, AP 2.6 mm posterior to bregma, DV 4.8 mm) and fixed to the skull using superglue (Loctite) and black dental acrylic (Ortho-Jet). Animals were allowed to recover for 3-4 weeks before experimentation.

### Fiber photometry from excitatory vlPAG neurons

In order to record neuronal activity from excitatory vlPAG neurons, 100 μL of *AAV9 CaMKII-jGCaMP8m-wPRE* (Addgene #176751) was injected into vlPAG of *Pnmt^2a-iCre^*;*Dbh^Flp^* mice and an optic fiber cannula was implanted above the injection site (ML -1.04 mm, 4.1 mm posterior from Lambda and DV 2.45 mm at 14° angle). The optic fiber cannula was fixed to the skull with superglue and black dental acrylic (Ortho-Jet). Animals were allowed to recover for 3-4 weeks before experimentation.

### DREADDs and fiber photometry

*Pnmt^2a-iCre^*;*Dbh^Flp^* mice were bilaterally injected with either *AAV8 hSyn-CONFON-HM3D-mCherry*^43^ (Addgene #183532) or *AAV8 hSyn-CONFON-HM4D-mCherry* (Fenno 2020, Addgene #177672) into RVLM. Next, the same mice were injected with 100 μL of *AAV9 CaMKII-jGCaMP8m-wPRE* (Addgene #176751) into the left vlPAG and an optic fiber cannula was implanted above the injection site, and fixed to the skull as described previously. Animals were allowed to recover for 3-4 weeks before experimentation.

### Immunohistochemistry and imaging

Mice were euthanized with tribromoethanol (240 mg/kg) and perfused transcardially with 10 mL PBS followed by 10 mL of 4% PFA. Brains were harvested, postfixed in 4% PFA overnight at 4 °C and cryopreserved in PBS containing 30% sucrose for 2-4 days at 4 °C. Brains were embedded in tissue freezing media (OCT) and kept at -80 °C until sectioning. 50 μm brain sections were made on a cryostat (CM1950, Leica Biosystems ) and collected into well plates filled with TBS (pH 8.4). Floating sections were washed for 10 minutes in TBS and incubated for 1 h in blocking solution (TBS, 5% NDS, 1% BSA, 0.5% Triton X-100). Sections were incubated overnight at 4 °C in antibody solution (TBS, 5% NDS, 1% BSA, and 0.3% triton 100) containing primary antibodies (goat anti-RFP (1:1000; Rockland #200-101-379), rabbit anti-GFP (1:1000; Invitrogen #A-11122), chicken anti-TH (1:250; Aves #AB10013440), rabbit anti-cFos (1:5000; Synaptic Systems #226008 or HelloBio #HB8006). Next, sections were washed twice for 10 minutes at room temperature with TBS and incubated for 90 minutes at room temperature with solution containing secondary antibodies (all at 1:500 dilutions; Cy3 donkey anti-goat (Jackson Immunoresearch laboratories #705-165-003); Cy3 donkey anti-rabbit (Jackson Immunoresearch laboratories # 711-165-152); 488 donkey anti-rabbit (Jackson #711-545-152); 647 donkey anti-chicken (Jackson Immunoresearch laboratories #703-605-155); 647 donkey anti-rabbit (Jackson Immunoresearch laboratories #711-605-152). Sections were then washed twice for 10 minutes at room temperature with TBS. Finally, sections were incubated for 10 minutes at room temperature with DAPI solution (Fisher, cat. #D1306, 1:25000 in TBS) and mounted with Fluoromount (Life Technologies cat. #00-4958-02). Representative image in 1c was obtained using a Zeiss LSM 700 confocal microscope and a 40X objective. Representative images in 4b were obtained using a X-Light V3 spinning disk confocal (CrestOptics) coupled to a Nikon Ti2 microscope and a 40X objective. Representative images in 2b, S1 and S5c were obtained using a Leica DM6 B epifluorescence microscope and a 20X objective.

### iDISCO tissue clearing

iDISCO+ tissue clearing method used as described previously with some modifications^104,105^. Briefly, mice were euthanized with an i.p. injection of tribromoethanol (Avertin) and were intracardially perfused first with 10 mL of cold PBS containing 0.02 mg/mL heparin, then with 10 mL 4% PFA. The brains were harvested and fixed overnight in 4% PFA with rotation at 4 ⁰C. All washes and incubations were performed with rotation unless otherwise specified. The samples were washed three times in PBS 30 minutes at room temperature followed by 1 h dehydration steps of methanol/H2O concentrations (20%, 40%, 60%, 80%), followed by two 1 h washes in 100 % methanol. Samples were incubated overnight in a mixture of dichloromethane (Millipore Sigma, cat. #270997) (DCM)/methanol (66% DCM and 33% methanol) at room temperature. Brains were then incubated in 100% DCM for 1 h, washed in 100% methanol for 1 h at room temperature and 1 h at 4 ⁰C. For the bleaching step, brains were incubated overnight in chilled 5% H2O2 (in methanol) at 4 ⁰C. Then, samples were rehydrated in 1 h steps using methanol/PTx2 (PBS, 0.2% Triton-X 100) and decreasing methanol concentrations (80%, 60%, 40%, 20%) at room temperature, followed by two 1 h washes in 100% PTx.2 at room temperature. Next, samples were incubated at 37 ⁰C for 72 h in permeabilization solution (PBS, 0.2% Triton-X 100, 20% DMSO, 0.3 M glycine). Then, samples were incubated at 37 ⁰C for 48 h in blocking buffer (PBS, 0.2% Triton-X 100, 10% DMSO, 6% NDS). After blocking, samples were incubated at 37 ⁰C for 7 days in PTwH (1 x PBS, 0.2% Tween-20, 10 μg/mL heparin, 6% DMSO, 3% NDS) containing primary antibodies rabbit anti-GFP (1:500) and goat anti-RFP (1:500). Next, samples were washed four times in 2X PTwH (10, 20, 30 and 60 minutes) and with 1X PTwH every 24 h for two days. Then the samples were incubated at 37 ⁰C for 7 days in PTwH containing secondary antibodies (1:500 donkey anti-rabbit Alexa fluor 647 and 1:500 Cy3 donkey anti-goat IgG (H+L) (Jackson Immunoresearch laboratories, cat. # 705-165-003). Then, samples were washed at room temperature for 3 days in PTwH. Next, samples were dehydrated with 1 h washes in a series of methanol/H2O concentrations (20%, 40%, 60%, 80%, 100%) and washed in 100% methanol overnight. The final delipidation step was done by incubating the samples in a DCM/methanol mixture (66% DCM and 33% methanol) for 3 h, then washed twice at room temperature in 100% DCM. Finally, samples were transferred to dibenzyl ether (DBE) for RI matching and left in the dark without rotation at room temperature for at least two days before imaging.

### Light Sheet Fluorescence Microscopy and image processing

All cleared brains were first imaged with an UltramicroscopeII dual illumination Gaussian light sheet instrument (Myltenyi Biotech) with pco.edge 4.2 M camera and an MVPLAPO 2X objective (Olympus) mounted on an MVX-10 zoom body with total magnification from .63-6.3x (Olympus). Illumination is driven by a laser module with four lines: 405 nm at 100 mW, 488 nm at 85 mW, 532 at 100 mW, and 639 at 70 mW. The samples were mounted ventral side up and imaged using Imspector Pro microscope controller software (v7). Images at horizontal plane acquired at 0.63x magnification with exposure time 100ms in z-stack at 5 μm intervals. Autofluorescence images were captured using 488 nm laser at 50 % laser power, RFP staining was imaged at 532 nm laser at 50 % laser power and GFP staining was imaged at 647 nm laser at 20 % laser power. High magnification images of rostroventral medulla (RVLM) was acquired with mesoSPIM open-source axially scanning light-sheet microscope with Photometrics Iris-15 camera, Oxxius L4Cc laser combiner and Mitutoyo Plan Apo Brightfield/Darkfield, 5X/0.14NA objective. Samples were mounted on a 3D printed adaptor to be optically sectioned through sagittal orientation and imaged using open-source mesoSPIM-control software. Images were acquired with exposure time 20 ms in z-stack at 2 μm intervals. RFP staining was imaged at 561nm at 50% laser power and GFP staining was imaged at 647 nm at 20% laser power. Images from both microscopes were processed in Arivis 4D software (version 4.1.2). GFP+ axons of the rabies tracing brains were segmented using TRAILMAP software^106^. Due to the high density of the axon labeling, RFP+ and GFP+ cells bodies in RVLM region of brains imaged at high magnification, were manually segmented using Arivis software (Zeiss). Spherical objects were placed on the GFP and RFP positive cell bodies by carefully navigating through each Z stack. The fluorescent positive cells were required to comprise of dendritic/axonal projections and basic neuronal soma anatomy to qualify for segmentation. Green spheres were used for GFP+ cell bodies and magenta spheres were used for RFP+ cell bodies.

### In situ hybridization plus immunohistochemistry

Mice were euthanized with an i.p. injection of tribromoethanol (Avertin) and were intracardially perfused with 10 mL of cold PBS and with 10 mL 4% PFA. Collected brains were postfixed overnight in 10% neutral buffered formalin, embedded in paraffin, sectioned at 4.5 μm in sagittal orientation, mounted on positively charged glass slides (Superfrost Plus; 12-550-15, Thermo Fisher Scientific) that were dried at 60 °C for 20 minutes, deparaffinized, and stained with hematoxylin and eosin (HE; Richard-Allan Scientific) or used for immunohistochemistry or in situ hybridization assays. HE-stained sections were covered using the HistoCore SPECTRA Workstain (Leica Biosystems). The RLVM was evaluated every 50 µm between interaural (lateral) coordinates 0.72 mm to 1.80 mm from the midline for each specimen^6^. All immunofluorescence (IF) staining was performed on Ventana Discovery Ultra Autostainer (Roche). All tissue sections used for immunofluorescence labeling underwent deparaffinization.

Primary antibodies were serially applied using the U DISCOVERY 5-Plex IF procedure (Ventana Medical Systems, Roche). Ready to use DISCOVERY OmniMap anti-Rb HRP (cat. #760-4311, 16 minutes), visualization of anti-RFP (Abcam, ab124754; 1:10,000, 64 minutes) was done using a DISCOVERY Rhodamine 6G kit (cat. #760-244, Roche). Detection of anti-GFP (Clontech, cat. #632381; Clone JL8, 1:2000, 64 minutes) was performed using a DISCOVERY RED610 kit (cat. #760-245, Roche). Anti-Thyrosine Hydroxylase (Millipore, AB152; 1:500, 64 minutes) was visualized using a DISCOVERY CY5 Kit (760-238). Discovery QD DAPI nuclear counterstain (cat. #760-4196, Roche) was applied. Finally, coverslips were mounted on slides with Prolong Gold Antifade reagent (Thermo Fisher Scientific, P36961).

A 4.5 µm section of brain from each mouse was also probed for RNAscope 2.5 VS Probe-Mm-Pnmt (cat. #426429; HRP Green, Advanced Cell Diagnostics) and RNAscope 2.5 VS Probe-Mm-Th-C2 (cat. #317629-C2; Fast Red, Advanced Cell Diagnostics) using a Ventana DISCOVERY ULTRA Autostainer (Ventana Medical Systems) and the RNAscope VS Duplex Reagent assay kits and the following products: mRNA RED Detection kit (Part number 760-234, 07099037001, Advanced Cell Diagnostics) and mRNA Teal Detection kit (Part number 760-256, Ordering code 08352941001, Advanced Cell Diagnostics). RNAscope 2.5 VS Duplex Control Probes (Ppib-C1, Polr2a-C2, Cat No. 320769, Advanced Cell Diagnostics)-Mm and RNAscope 2.5 VS Duplex Negative Control Probe (DapB-C1, DapB-C2, Cat No. 320759, Advanced Cell Diagnostics) were used to assess specimen quality and the immunolabeling procedure in serial tissue sections. Subsequently, immunohistochemistry (IHC) was performed with anti-GFP (Clontech, #632381; Clone JL8, 1:2000, 32 minutes) using heat-induced epitope retrieval with cell conditioning media 1 (CC1, 950-224, Ventana Medical Systems) for 32 minutes at 37 °C followed by visualization with DISCOVERY OmniMap anti-Rb HRP (760-4311; Ventana Medical Systems), and the DISCOVERY ChromoMap DAB kit (760-159; Ventana Medical Systems). Positive (adrenal gland) and negative (lymph nodes and liver) tissue controls and single ISH and IHC were used to assess the specificity and sensitivity of immunostaining of markers used in any multiplex panels. Sections were digitized with a Zeiss Axiovision Scanner (Zeiss) and analyzed using HALO v3.6.4134.137 for serial image registration of the serial IF and ISH/IHC tissue sections. IF and ISH staining was quantified using the ISH-IHC v2.2 algorithm.

### Plethysmography and respiratory analysis

Plethysmography was performed in 14–20-week-old mice as previously described^107^. Briefly, mice were individually placed in a 450 mL whole animal plethysmography chamber (EMKA technologies) at room temperature (22°C) and allowed to acclimate for at least 30 minutes or until exploratory movements stopped. Breathing was then recorded, and inspiratory and expiratory time, and breathing frequency were determined using the EMKA iOX2 software.

### Behavioral experiments

For all behavioral experiments, infrared light sources were used, and a video camera (White Matter e3Vision or Basler acA2040-90um-NIR)was located overhead to record the activity of the animals. Mouse tracking was obtained from videos using DeepLabCut^108,109^ and outputs were analyzed with R and DLC-ROI software.

### Open Field Test (OFT)

The open field test was used to assess anxiety-like behaviors in mice as previously described^110^. Briefly, mice were transferred into the testing room to acclimate for at least 30 minutes or until there was a large reduction in exploratory behaviors. Mice were placed in the center of the open field (OF; 40 x 40 x 20 cm) and allowed to freely explore the arena for 20 minutes. No acclimation to the OF was done prior to testing. Distance traveled, velocity, and time spent in the center were quantified.

### Elevated zero maze (EZM)

To measure anxiety-like behaviors using the EZM^60,111^, mice were transferred into the testing room and allowed to acclimate at least 30 minutes or until there was a large reduction in exploratory behaviors. The EZM consisted of a circular maze (100 cm diameter, lane width 10 cm and elevated 61 cm from the floor) with 2 out of 4 quadrants enclosed with 20 cm walls. Mice were placed in one of the open areas of the EZM and allowed to freely explore for 20 minutes. One acclimation trial was performed one day before testing. Distance traveled, velocity and time spent in the open and closed areas of the maze were quantified.

To test the chronic effects of stimulation of C1 neurons, mice were transferred into the testing room and allowed to acclimate at least 30 minutes or until there was a large reduction in exploratory behaviors. Next, each mouse received optogenetic stimulation in their home cage. The mice were immediately transferred back to the housing room and placed into the rack. Mice remained undisturbed for seven days, after which they were transferred into the testing room for acclimation as described above. Then, mice were placed on the EZM and allowed to explore freely for 20 minutes. For this testing, an acclimation trial was omitted to minimize handling of the mice.

In experiments involving the use of DREADDs, an i.p. injection of deschloroclozapine (DCZ, Fisher #71-931-0, 500 μg/kg) or saline solution (0.9 % NaCl) was delivered 15 minutes before testing. In experiments involving physical restraint, mice were placed in a homemade restrainer (built using a 50 mL falcon tube with holes drilled on the cone to allow sufficient airflow) for 30 minutes. Mice were then released directly into one of the open areas of the EZM. In experiments involving DREADDs and restraint, mice were given an i.p. injection of DCZ (500 μg/kg) and immediately placed inside the restrainer for 30 minutes before testing.

### Optogenetic stimulation

In experiments involving optogenetic stimulation, an optic fiber patch cord was connected to the mice, and they were released into open areas of the EZM or in the center of the OFT. Optogenetic stimulation was carried out with an Optogenetics-LED box (Prizmatix), a patch cord attached to a low-AF rotary joint and a patch cord to ferrule of 0.67 N.A. The stimulation parameters were as follows: 20 Hz, pulse width 20 ms, power ∼5.5 mW (measured at the tip of the patch cord). The stimulation train consisted of 30 second stimulation followed by 30 seconds rest, for a total time of 10 minutes.

### Real-Time Place Preference (RTPP)

Real-time place preference was performed as previously described^66^. Briefly, an arena (40 x 40 x 30 cm) was divided in two halves (stimulation and non-stimulation sides) by a 10 cm tall plastic wall. An aperture of 6 cm wide communicates both sides of the chamber. A tracking system was used to automatically trigger the optogenetic LEDs. Mice were placed on the non-stimulation side and received optogenetic stimulation upon crossing to the stimulation side (20 Hz, 20 ms). Stimulation stopped automatically when mice returned to the non-stimulating side. The total duration of the test was 20 minutes.

### Fiber photometry

Fiber photometry was performed using a LUX RZ10X and Synapse software (Tucker-Davies Technologies). Excitation light was provided by 405 nm (Isosbestic, Lx405) and 465 nm (Calcium, Lx465) LEDs. The signals were acquired using 2^nd^ generation photosensors (LxPS2) at 1 kHz. Low-autofluorescence optic fiber patch cords were used (Doric lenses, cat. #MFP_400/430/1100-0.57_1m_FC-ZF1.25 (F)_LAF). A low-autofluorescence rotary joint was used to prevent entanglement of the patch cord (Doric Lenses, cat. #FRJ_1x1_PT-G2_400-0.57_1m_FCM_0.15m_FCM). Synchronization between camera feed and fiber photometry recordings was performed in real time using a PC running Bonsai software and an Arduino board to trigger a signal when mice entered an ROI. Fiber photometry data was analyzed using Guided Photometry Analysis in Python (GuPPy)^112^.

### Statistics and reproducibility

No statistical methods were used to predetermine sample size. Data were excluded in certain conditions (missed viral injections for behavioral experiments). Data distribution was assumed to be normal, but this was not formally tested. Investigators were not blinded to mouse genotype during experiments or quantification of data. However, care was taken to ensure that a similar number of male and female mice were included across genotypes and experimental conditions.

## Acknowledgements

We thank Danielle Dunwald and Hunter Nolen for technical support, Dr. Steven Ebert (University of Central Florida) for sharing the *Pnmt^Cre^* mouse line, the St. Jude Vector Core Lab for generating AAVs, the St. Jude Center for Advanced Genome Engineering (CAGE) and Valerie Stewart (Director, St. Jude DNB Neuroembryology Core) for generation of the *Pnmt^2a-iCre^* transgenic mouse line, Heather Sheppard from the St. Jude Comparative Pathology Core for assistance with *in situ* and immunostaining multiplex experiments, Daniel Stabley, Sharon King and Abbas Shirinifard from the St. Jude DNB Neuroimaging Core for assistance with light sheet imaging and analysis, and members of the L.A.S. laboratory for helpful feedback. This work was supported by a NARSAD Young Investigator Grant from the Brain & Behavior Research Foundation (to C.F.P.), the NIH (grant no. 1DP2NS115764 to L.A.S.), and institutional funds from St. Jude Children’s Research Hospital (to C.F.P., L.M.F., R.L.P., B.G.P., and L.A.S.).

## Author contributions

C.F.P. and L.A.S. conceived the project. C.F.P. designed and performed experiments and analyzed data. R.L.P. performed stereotaxic surgeries and histology, L.M.F. performed brain clearing experiments and B.G.P. performed histology. L.A.S. generated viruses, designed the *Pnmt^2a-iCre^*mouse line, and supervised the project. C.F.P. and L.A.S. wrote and edited the paper with feedback from the other authors.

---

**Extended Data Figure 1.**
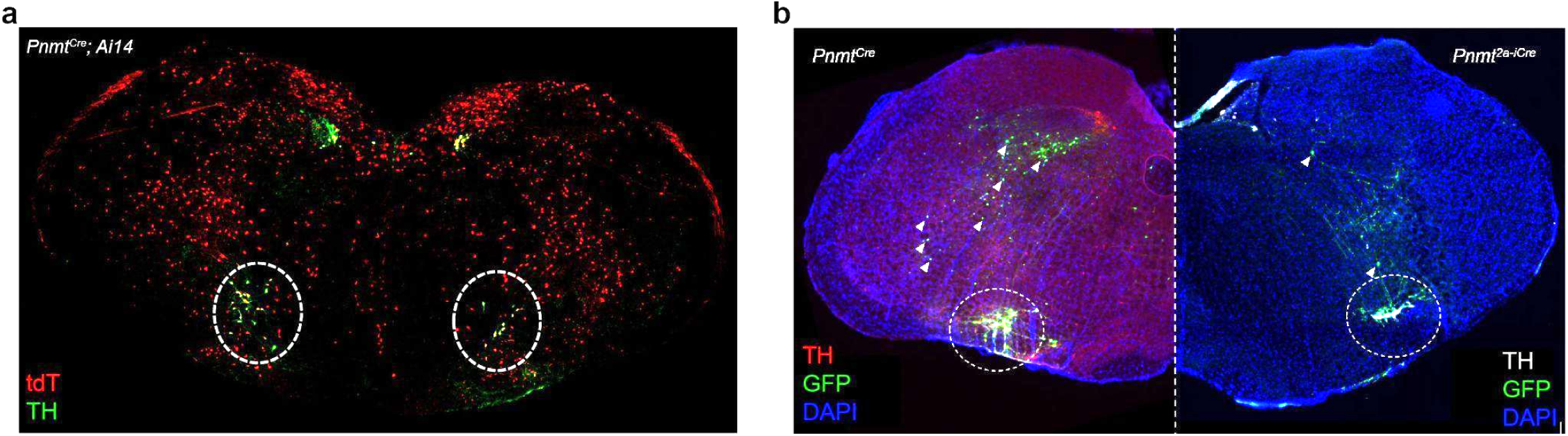
Comparison of Cre expression patterns between transgenic lines targeted to the *Pnmt* locus. **a,** Representative image from a *Pnmt^Cre^;Ai14* animal shows widespread ectopic expression of the Cre-dependent reporter tdTomato (red) beyond defined TH^+^ catecholaminergic cells (green) in the rostroventrolateral medulla (RVLM, dotted circles). n=3 animals. **b,** Representative images comparing expression of Cre-dependent AAV-GFP (green) injected into RVLM (dotted circles) in *Pnmt^Cre^* (left) and *Pnmt^2a-iCre^* (right) transgenic animals. Ectopic expression beyond RVLM catecholaminergic cells (TH, red) is greatly reduced in *Pnmt^2a-iCre^* tissue (white arrowheads), suggesting improved restriction of Cre expression in these animals. Nuclei are counterstained with DAPI (blue). n=3 animals of each genotype.

**Extended Data Figure 2.**
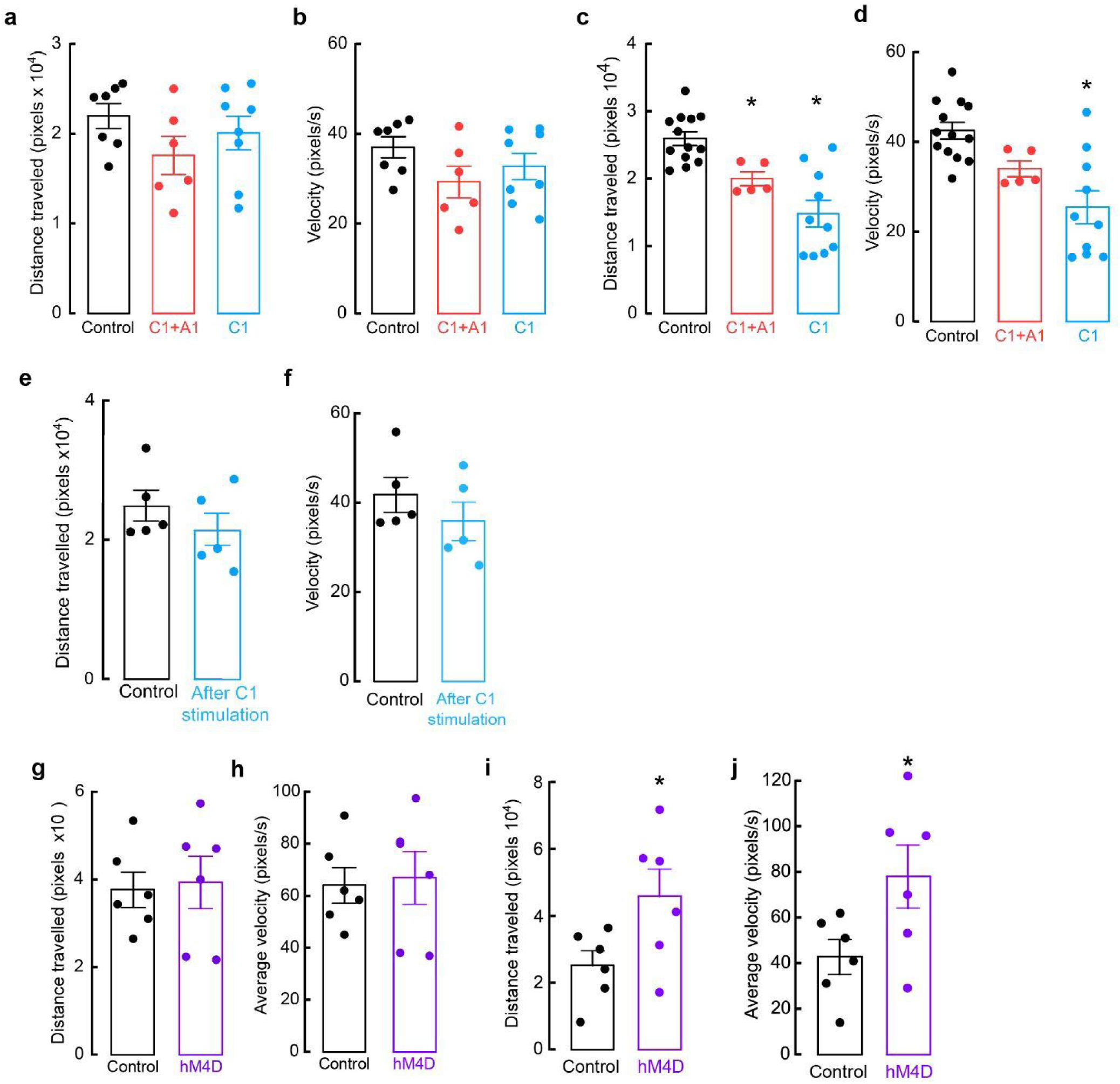
Assessment of distance traveled and velocity during EZM in animals undergoing optogenetic or chemogenetic manipulation. **a,** Average distance traveled and **b,** average velocity of control (black), C1+A1 (red) and C1 (blue) mice with optic fiber implantation in RVLM. Control n=7, C1+A1 n= 6, C1 n=8. **c,** Average distance traveled and **d,** average velocity of control (black), C1+A1 (red) and C1 (blue) mice with optic fiber implantation in PAG. Control n=13, C1+A1 n=5, C1 n=10. *p<0.05 vs control, Kruskal Wallis test with Dunn’s post hoc test. **e,** Average distance traveled and **f,** average velocity of control (black) and C1 (blue) mice with optic fiber implantation in PAG. Control n=5, C1 n=5. Testing occurred approximately one week after mice received optogenetic stimulation in their homecage (20 Hz, 20 ms pulse width, 30 s stimulation, 30 s rest, for 10 min total). **g,** Average distance traveled and **h,** average velocity of control (black) and C1 hM4D (purple) mice after administration of the DREADD agonist DCZ. Control n=6, hM4D n=6. **i,** Average distance traveled and **j,** average velocity of control (black) and C1 hM4D (purple) mice after administration of vehicle or DREADD agonist DCZ and prior exposure to restraint. Vehicle n=4, DCZ n=4. Data represents mean ± SEM. *p<0.05, Mann-Whitney U test.

**Extended Data Figure 3.**
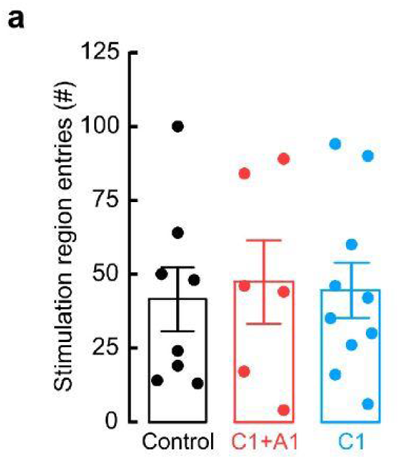
Optogenetic activation of A1 or C1 axons innervating vlPAG does not alter entries to the stimulation side in the RTPP test. **a,** Average number of entries to the stimulation side of the chamber in the RTPP during the duration of the test (20 min) of control (black), C1+A1 (red) and C1 (blue) mice. Control n=8, C1+A1 n=6, C1 n=10. Data represents mean ± SEM.

**Extended Data Figure 4.**
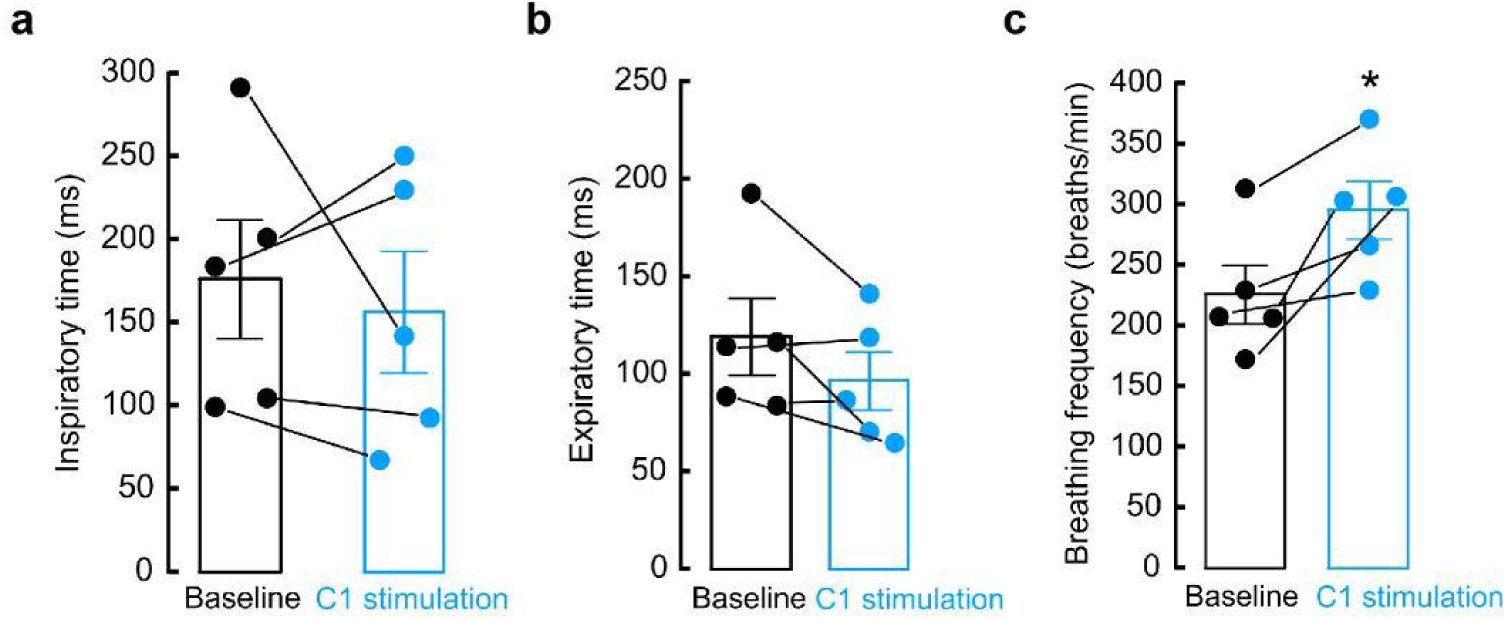
Optogenetic activation of C1 axons innervating vlPAG increases breathing rate without altering inspiratory and expiratory duration. **a,** Average inspiratory duration, **b,** expiratory duration, and **c,** breathing frequency of mice before (baseline, black) and during optogenetic stimulation (C1 stimulation, blue). Data represents mean ± SEM. *p<0.05, Mann-Whitney U test.

**Extended Data Figure 5.**
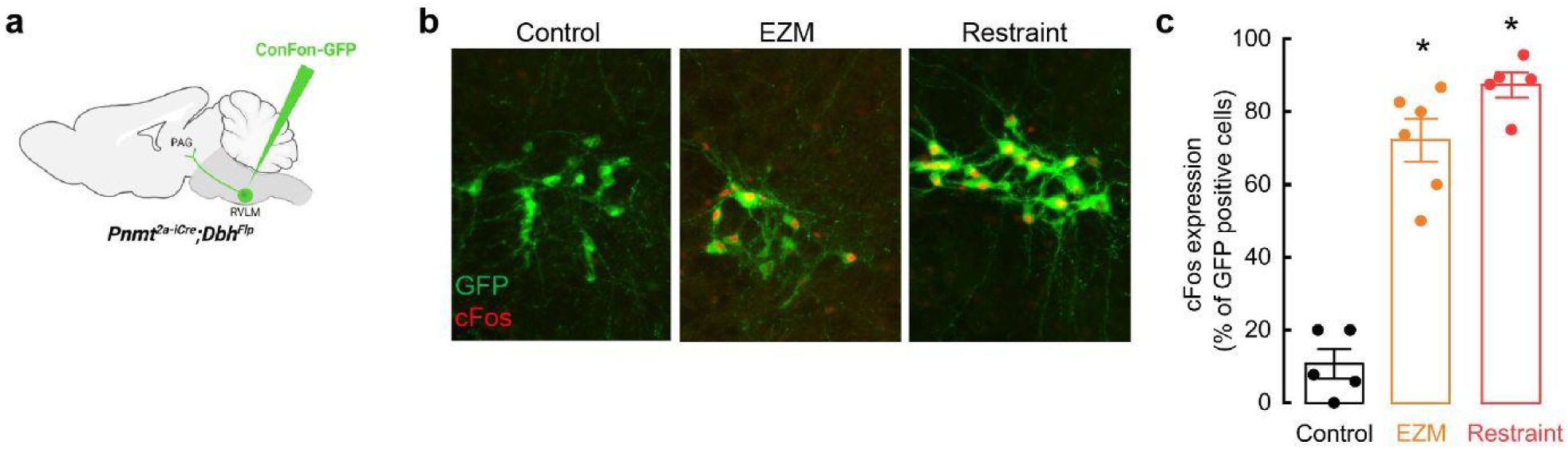
Stressful stimuli activate C1 neurons. **a,** Strategy for intersectional viral expression of GFP to label C1 neurons. **b,** Representative images showing increased expression of cFos (red) in C1 neurons (green) following stressful experiences (EZM and restraint). **c,** Average percentage of GFP-labeled C1 neurons that express cFos in control (n=5), EZM (n=6) and restraint (n=5) conditions. Data represents mean ± SEM. *p<0.05 vs control, Kruskal Wallis test with Dunn’s post hoc test.

**Extended Data Figure 6.**
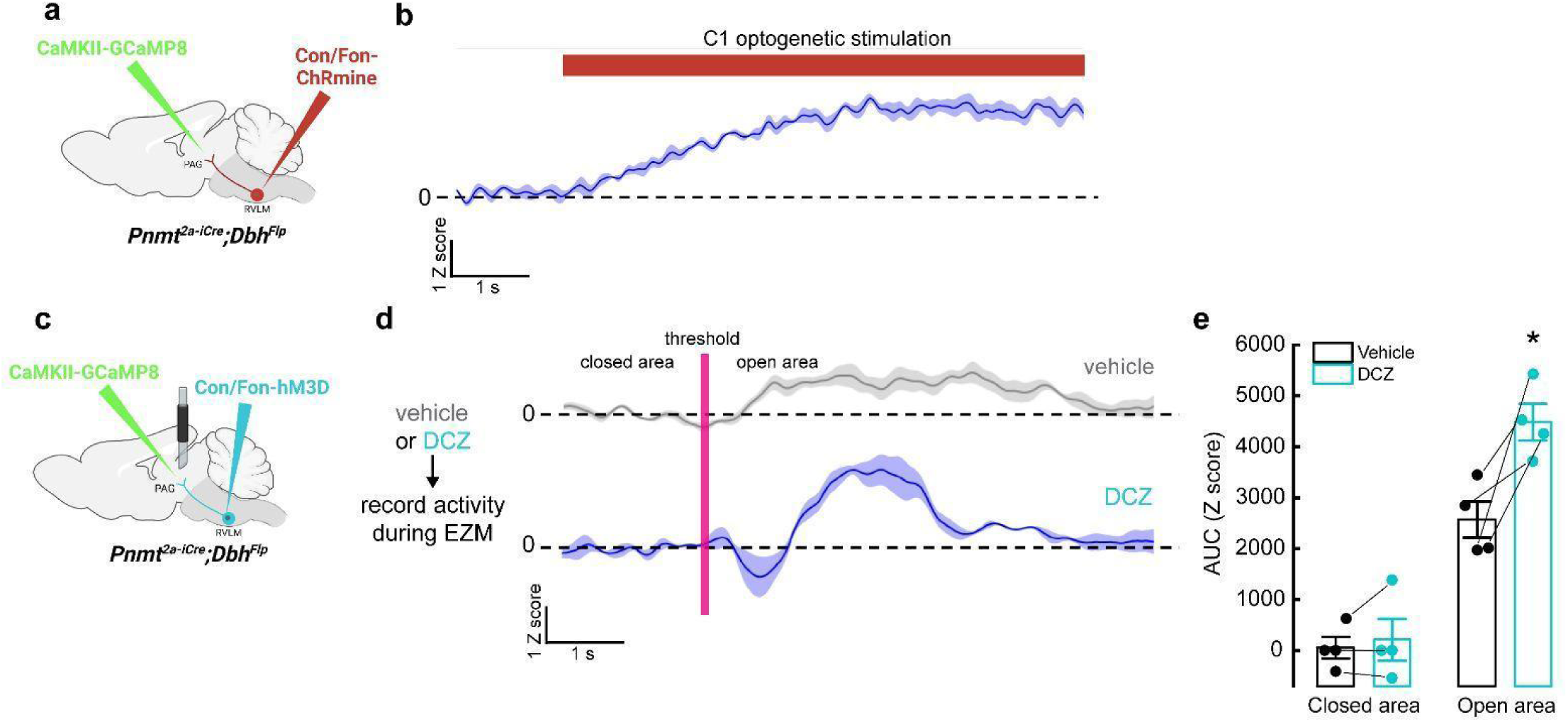
Optogenetic and chemogenetic activation of C1 neurons increases excitatory neuron activity in vlPAG. **a,** Schematic of the approach for optogenetic activation of C1 neurons while recording bulk calcium responses from excitatory vlPAG neurons. **b,** Representative average calcium activity of excitatory vlPAG neurons to optogenetic stimulation of C1 neurons (red line, 20 Hz, pulse duration 20 ms, total duration 10 s) under anesthesia. Data represents the mean Z score (dark line) ± SEM (light color) of 3 trials. **c,** Schematic of the approach for chemogenetic activation of C1 neurons while recording bulk calcium responses from excitatory vlPAG neurons. **d,** Representative average calcium activity of excitatory vlPAG neurons while mice transition from the closed to the open area of the EZM, in mice pretreated with vehicle (grey) or DCZ (purple). Data represents the mean Z score (dark line) ± SEM (light line) of 3 trials. **e,** Average AUC analysis of excitatory vlPAG neuron calcium activity from -5 to 0 seconds before crossing the threshold (closed) and from 0 to 5 seconds after entering the open area (open). Data represents mean ± SEM. n=4 animals. *p<0.05, Mann-Whitney U test.

**Extended Data Figure 7.**
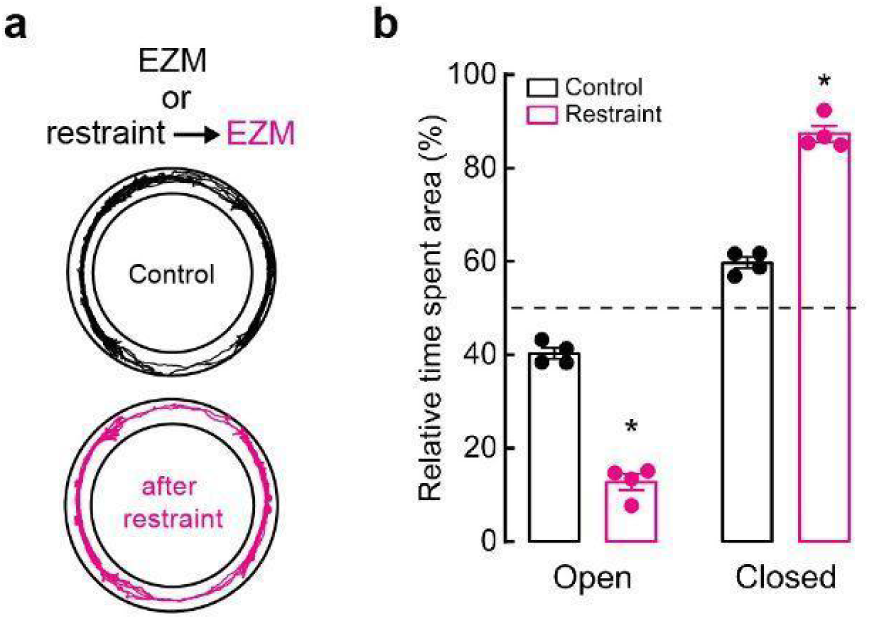
Pre-exposure to restraint enhances anxiety-like behavior in EZM. **a,** Representative cumulative trajectories (10 min) in EZM testing for without (Control, black) or with pre-exposure to restraint (30min)(pink). **b,** Average percentage of time spent in the open and closed areas of the EZM. Control n= 4, Restraint n= 4. Data represents mean ± SEM. *p<0.05, Mann-Whitney U test.

